# A simple approach for image-based modelling of the heart that enables robust simulation of highly heterogeneous electrical excitation

**DOI:** 10.1101/2022.11.11.516084

**Authors:** Michael A Colman, Alan P Benson

## Abstract

Remodelling of cardiac tissue structure, including intercellular electrical coupling, is a major determinant of the complex and heterogeneous excitation patterns associated with cardiac arrhythmias. Evaluation of the precise mechanisms by which local tissue structure determines global arrhythmic excitation patterns is a major challenge that may be critically important for the development of effective treatment strategies. Computational modelling is a key tool in the study of cardiac arrhythmias, yet the established approaches for organ-scale modelling are unsuitable to capture the impact of local conduction heterogeneities; a novel approach is required to provide this multi-scale mechanistic insight.

We present a fundamentally simple yet powerful approach to simulate electrical excitation in highly heterogeneous whole-heart models that exploits the underlying discreteness of the myocardium. Preliminary simulations demonstrate that this approach can capture lower conduction velocities and reproduce wave breakdown and the development of re-entry in conditions where the established approaches cannot.

## 1 Introduction

Cardiac arrhythmias are a major cause of morbidity and mortality associated with cardiovascular disease (CVD). Loss of the regular rhythm of the heart can substantially reduce cardiac output, detrimentally affecting the delivery of oxygen and nutrients to the vital organs of the body. Acute arrhythmia events can be fatal and, indeed, sudden cardiac death accounts for a substantial proportion of mortalities associated with CVD^1^.

Electrical and structural remodelling occur over the progression of multiple CVDs and it is well established that this generally promotes both the initiation and sustenance of arrhythmia^2,3^. Recent studies have highlighted the mechanistic role by which structural remodelling promotes arrhythmia: conduction heterogeneity contributes to unidirectional conduction block and enables the development of sustained re-entrant circuits^4,5^. However, due to the conflict in spatial scales required to simultaneously resolve local conduction while maintaining the global picture of whole-heart activation, it is a major challenge to fully dissect and elucidate the precise mechanisms by which intercellular coupling and microstructure determine arrhythmia dynamics.

Multi-scale computational modelling has proved an invaluable tool for elucidation of the complex mechanisms underlying arrhythmia; such insight can help to drive novel pharmacological and surgical interventions to prevent, manage, or treat CVD in a diverse population^6–8^. The two most commonly used numerical methods for simulation of organ-scale cardiac electrophysiology are the finite difference method (FDM) and the finite element method (FEM), referring to different mathematical approaches to discretise the mono- or bi-domain reaction-diffusion equations which describe spatial diffusion of electrical or chemical activity^9^. Both of these approaches, however, are limited in their ability to capture the discrete underlying details of electrical coupling between myocytes, and are furthermore not well-suited to modelling the impact of locally heterogeneous conduction associated with structural remodelling, including the proliferation of fibrosis.

Recently, cellular-scale tissue models have been developed which describe intercellular coupling to a much greater level of sophistication^10–13^, accounting for the influence of intra- and extracellular spaces, dynamic gap-junction conductance, and the potential role of ephaptic coupling. However, large-scale, whole-chamber image-based models of the heart are computationally intractable to perform at cellular-scale resolutions, and it is not trivial to extend these approaches to the whole-heart scale. There is therefore a strong motivation to develop an approach to discretise image-based tissue models that readily facilitates highly heterogeneous conduction properties and is compatible with these more sophisticated models of electrical coupling. Such a model would offer the possibility to reveal new insight into the mechanisms of cardiac (dys)function and develop ever more accurate models of the heart, including for genuine subject specificity (for example, in patient specific clinical models). Here, we present a fundamentally simple yet powerful network model to achieve this goal. This manuscript will discuss the derivation of the approach and perform demonstration simulations that illustrate how it may be used to model heterogeneous conduction conditions.

## 2 Results

### 2.1 Connection maps in 2D and 3D

The presented approach creates a network of axial and transverse inter-nodal connections which are weighted based on local myocyte orientation on a 2D or 3D structured grid tissue geometry (**Fig. 1a,c**). These weightings determine the magnitude of the nodal conductance values that contribute to the junctional conductance value at each network connection (**Fig. 1b**). Weighted network connections are illustrated for three idealised cases (with different, global orientation directions) in a 2D sheet (**Fig. 2a - left**). It is worth noting that when the orientation points either exactly along the axis or diagonal, the junctional conductance in that direction is exactly equal to the axial conductance (*g*_a_) and the transverse coupling is applied also only in one direction (perpendicular); in these special cases, only two directions have a non-zero weighting (one axial and one transverse; **Fig 2a**).

**Figure 1.**
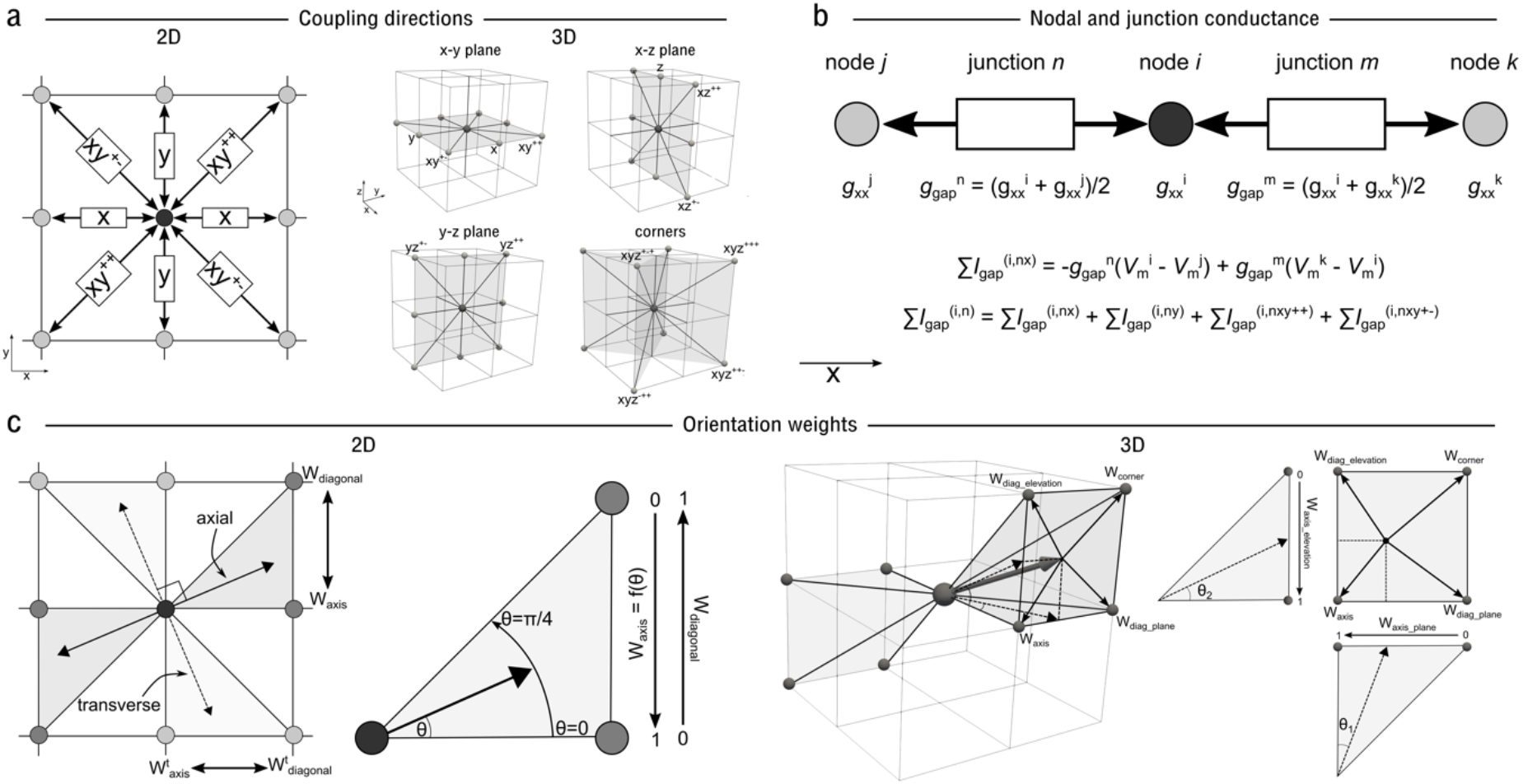
Summary of model approach. **a** – illustration of the different coupling directions and terminology referring to these directions, in 2D (left) and 3D (right). **b** - illustration of how the nodal and junctional conductance parameters are related and contribute to the total gap junctional conductance, for three nodes i, j, and k and two junctions n and m. **c** - illustration of the relationship between myocyte orientation angle and the weighting terms towards the axis and diagonal within a segment (2D) and towards the axis, in-plane diagonal, elevation diagonal, and corner in a quadrant (3D).

**Figure 2.**
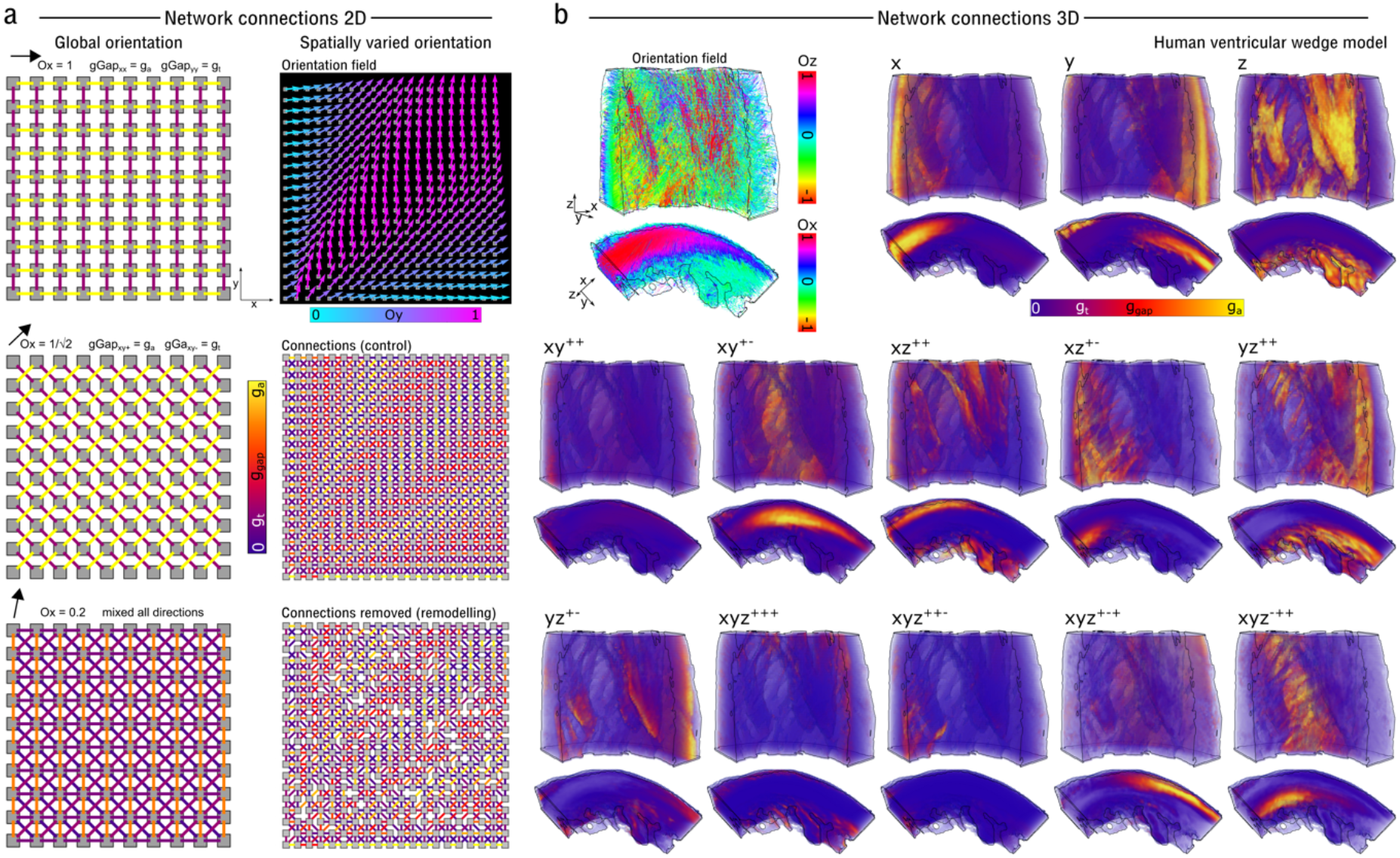
Illustration of weighted network connection maps. **a** - illustration of the connection maps in 2D for idealised cases with a global myocyte orientation in three different directions (left) and in the case of a spatially varying orientation field (right) for both control (g_a_ and g_t_ are spatially homogeneous; upper) and in a case where 20% of axial and 80% of transverse connections have been randomly removed (lower). Global orientation maps are illustrated on a 10×10 grid for clarity in visualisation. **b** - Illustration of directional condutances in a 3D human ventricular wedge model. Myocyte orientation streamlines are shown to provide context, with colour corresponding to either the z- or x-component (dependent on the view), along with the magnitude of the connection for each direction. The anisotropy ratio is 4:1 (i.e, g_a_ = 4g_t_). Note that, for clarity of visualisation, the junctional conductances defined in equations (37–39) have not been scaled by the 1/Δx etc factors, in order to normalise between axis and diagonal conductances.

Inter-nodal connections are further illustrated in 2D for the more complex condition with a spatially varying myocyte orientation vector field (**Fig. 2a - right**) in both control (all connections preserved) and remodelling (a set proportion of axial and transverse cellular connections have been removed.) It is worth explaining the patterns that are observed in the cellular connections, for clarity of interpretation of how the model works. If the orientation points exactly in x, the x-component of the nodal conductance, *g*_xx_^node^, will be exactly equal to *g*_a_ (the axial weight, *W_xx_* = 1, and transverse weight, *W*_xx_^t^ = 0), *g*_yy_^node^ will be exactly equal to *g*_t_ (*W*_yy_ = 0; *W*_yy_^t^ = 1) and there will be no diagonal connections (*W*_xy++_ = *W*_xy+-_ = 0). As the orientation is rotated towards the diagonal, the axial weight for the x direction will linearly reduce from 1 to 0 (while the weight for the diagonal linearly increases from 0 to 1) and the transverse weight for the y direction also linearly reduces from 1 to 0. Thus, exactly at the diagonal, *g*_xx_^node^ and *g*_yy_^node^ are 0 for both axial and transverse. As the orientation is rotated further towards y, the transverse weight for x increases from 0 to 1 along with the axial weight for y. Thus, *g*_xx_^node^ is smallest when the orientation points towards the diagonal, increases to g_t_ as the orientation rotates towards y, and increases to g_a_ as the orientation rotates towards x.

Connection maps are shown for two 3D geometrical models: a human ventricular wedge^14^ where only the myocyte orientation was given (and so the two transverse directions were calculated within the model; **Fig. 2b**), and a full reconstruction of rat bi-ventricular geometry^8^ wherein all three orientations describing cardiac tissue structure (myocyte, sheet, and sheet normal, corresponding to the three DT-MRI eigenvectors) were given (Online Supplement **Fig. S5**).

### 2.2 Verification of conduction patterns and comparison to FDM

The network model was parameterised to match FDM implementations regarding axial and transverse conduction velocity (see “Methods: Cellular dynamics and model parameters”). There were some notable differences in the conduction patterns in 2D and 3D between the two implementations (**Fig. 3**), but the overall patterns and activation time match well, and it is not necessarily the case that FDM represents a “ground truth”. Note that the spatial pattern in both methods when the orientation is diagonal is not an identical symmetrically rotated version of the pattern when the orientation is in-axis. This is discussed further in the **Online Supplement**.

**Figure 3.**
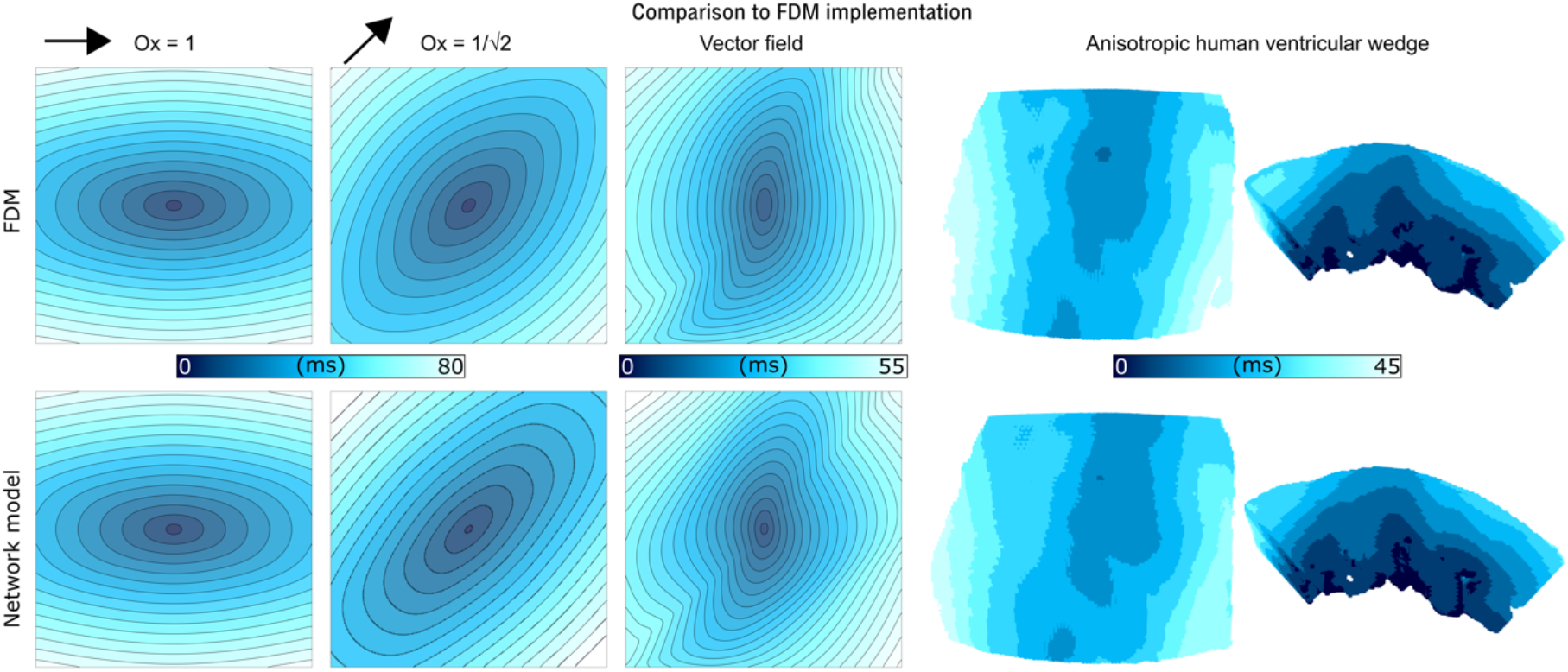
Comparison between activation patterns in parameter matched FDM and network model simulations in 2D with different global orientations (fully in x, exactly in xy^++^), and in the heterogeneous vector field shown in Fig 2, as well as in the 3D ventricular wedge model. Simulations with global orientation direction were performed on a 300×300 grid at a spatial resolution of Δx = 0.25 mm with coupling parameters of D1 = 0.4 mm/ms and D2 = 0.1 mm/ms (FDM method) and g_a_ = 1.6 nS/pF and g_t_ = 0.4 nS/pF (network model). Simulations using the vector field for myocyte orientation were performed on a 200×200 grid at a spatial resolution of Δx = 0.2 mm with coupling parameters of D1 = 0.2 mm/ms and D2 = 0.05 mm/ms (FDM method) and g_a_ = 0.8 nS/pF and g_t_ = 0.2 nS/pF (network model). Simulations on the 3D human ventricular wedge were performed at a spatial resolution of Δx = Δy = 0.2125 mm and Δz = 0.25 mm with coupling parameters of D1 = 0.1171 mm/ms and D2 = 0.0130 mm/ms (FDM method) and g_a_= 0.55nS/pF and g_t_ = 0.061 nS/pF (network model).

### 2.3 Conduction patterns in heterogeneous media

The impact of removing cellular connections at defined probability thresholds was demonstrated using a 2D model, implementing the heterogeneous vector field illustrated in **Fig. 2a**, across total spatial extents of 50’50 nodes and 200×200 nodes, enabling direct relation to visualised connection maps (**Fig. 4** and **Supplementary Videos 1 and 2**). Removing axial and transverse connections caused different spatial patterns of activation to emerge which clearly corresponded to the differential loss of connections in these directions. Non-uniformity in the propagating wavefront was observed in all conditions where cellular connections were removed. Patterns in media with sampled distributions of scale factors (rather than the binary choice of preserved/removed) are shown in the **Online Supplement Fig. S6**.

**Figure 4.**
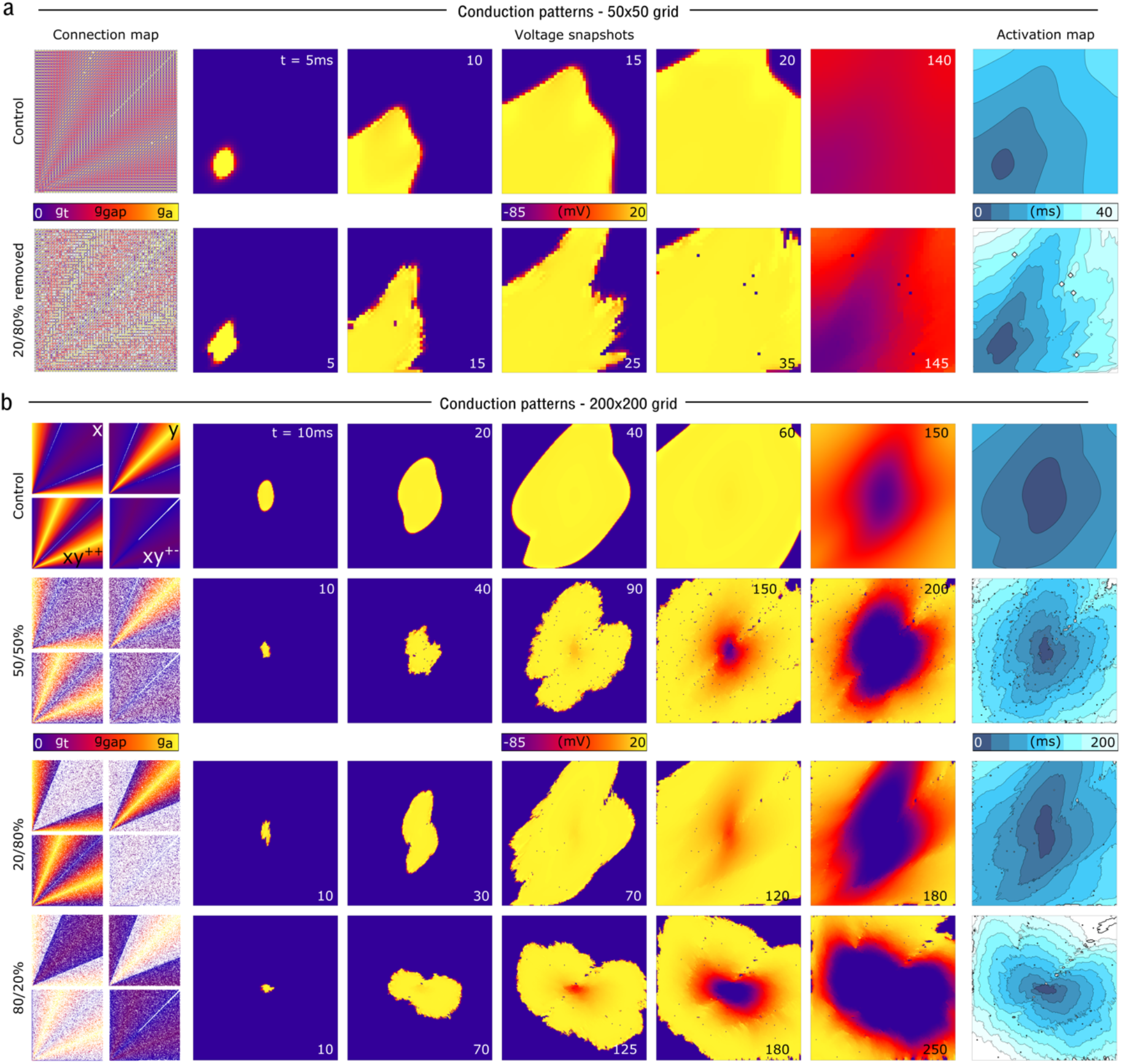
Conduction patterns in different connection-removal conditions. **a** – Connection maps and activation patterns in a 50×50 grid, where the pattern can be somewhat directly visually related to the connection maps. Control refers to the condition where all connections are present, and the bottom row is the condition where 20% of the axial and 80% of the transverse connections are randomly removed. **b** – connection maps and conduction patterns in four different conditions (control and cases where 50%-50%, 20%-80% and 80%-20% axial-transverse connections have been removed) in a 200×200 grid, where direct visual relation to the connection map is more difficult, but the tissue has sufficient area to permit a reasonable activation time.

Preliminary simulations implementing heterogeneously removed cellular connections revealed that these conditions were sufficient to promote wavefront breakup (**Fig. 5a**), which, in combination with rapid pacing, could lead to the development of re-entry (not observed with a parameter-matched global reduction to intercellular coupling in FDM; **Fig. 5b** and **Supplementary Videos 3 and 4**). It should be clarified that the regimes shown in these cases correspond to a substantial loss of connections and thus slow global conduction velocities and activation times, and this is reflected in square-like waves in the FDM implementation.

**Figure 5.**
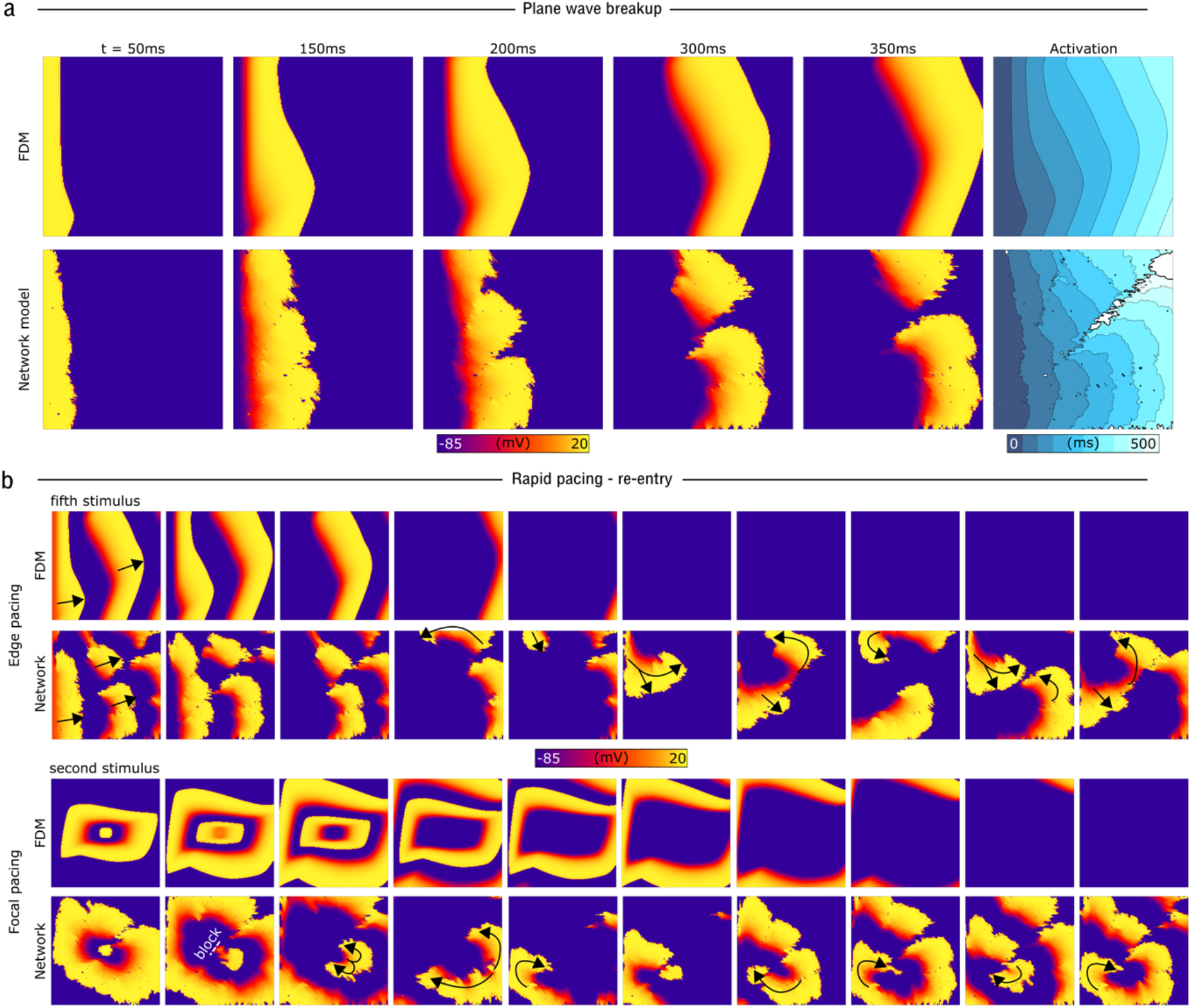
Illustration of wave-break and re-entry occurring in the network model with reduced connections. **a** – Voltage snapshots and activation pattern in the 2D model with the same vector orientation field as previous figures, paced from one edge of the tissue. **b** – Voltage snapshots in two different pacing scenarios (edge pacing for 5 stimuli, upper, and centre focal pacing for 2 stimuli, lower) on the same model as **a**. Sites of conduction block and wave propagation direction are labelled for clarity.

## 3 Discussion

### 3.1 Summary

We have presented a novel approach to construct a network model to describe cardiac intercellular electrical coupling on structured-grid geometries. The approach calculates coupling weights between adjacent nodes based on the local myocyte orientation (**Figs. 1–2**). We have demonstrated that this approach leads to expected conduction patterns in idealised and non-idealised tissue (**Figs. 3–5**). We have further demonstrated that this approach readily facilitates the direct manipulation of intercellular connections (**Fig. 2**), enabling complex conduction patterns to be simulated, such as observed with high levels of fibrosis. The model conserves current, is highly stable to complex media, and naturally accounts for boundary conditions.

### 3.2 Value of the model

Of the established approaches, FDM is the simplest but has limits regarding numerical stability in highly heterogeneous or anisotropic conditions; FEM is more complex and robust but is similarly not well-suited to modelling interrupted cellular connections. Neither model satisfactorily reproduces very slow conducting re-entrant circuits^15^. One possible explanation for why these established methods are unsuitable for simulation of these features is that they are derived from a mathematical discretisation of a spatially continuous system of equations, yet cardiac tissue is inherently discrete at a sufficiently large spatial scale (that of the myocyte: 10-100 μm) that this discreteness may be important for dynamics. Our new approach exploits this underlying discreteness in its fundamental philosophy, presenting advantages over these previous methods in computational efficiency, simplicity, numerical stability (and consequent ability to solve with simple numerical methods), and mass conservation. However, the main advantages, and primary motivation for the development of the model, are as follows:

i. the ability to directly modify or remove cellular connections, resulting in representation of barriers to conduction between excitable regions.
ii. compatibility with the more sophisticated models of intercellular electrical coupling^10,12,13,16^, including models of dynamic, voltage-gated gap-junction conductance.

Implementation of the model readily enables direct replacement of previous implementations using FDM (the same geometrical and myocyte orientation reconstructions can be used, and conduction parameters easily matched). For this purpose, the model is made available open-source in two different packages: i) packaged with the Multi-scale cardiac simulation framework^17^ and ii) in a simplified, single-file implementation in C++ to be extracted and used in any context. Tools to generate heterogeneous connection maps are also provided with the code, all available at http://physicsoftheheart.com/downloads.html and https://github.com/michaelcolman/.

### 3.3 Comparison to previous models of fibrosis

Previous tissue simulations have typically represented fibrosis through a reduction of the diffusion coefficient and increase in the anisotropy ratio^18,19^, reflecting the slower overall conduction velocities and longer activation times associated with fibrosis and structural remodelling. This is generally applied homogeneously (i.e., the diffusion coefficient is reduced globally in all fibrosis regions) but may be applied according to spatial gradient maps^19^. Our approach does not rely on changes to the coupling strength, but instead reproduces reduced coupling by removing individual intercellular connections. A change in the anisotropy ratio consequentially arises if axial and transverse connections were differentially removed (**Fig. 4**).

Example simulations demonstrated that removing cellular connections can produce complex, heterogeneous, and highly anisotropic conduction patterns which may degenerate into arrhythmic dynamics; longer activation times and slower average conduction velocities were reproduced but with a distinct underlying pattern to a simple global reduction in intercellular coupling (**Fig. 5**). The importance of this feature is highlighted by considering conduction patterns in low coupling regimes: the FDM method produces unphysiological square-like waves, whereas the new approach captures the same overall reduction in activation time but with highly heterogeneous underlying wavefronts. Importantly, re-entry could emerge from regular rapid pacing, rather than requiring a well-timed S1-S2 stimulus or other complex protocol, more closely reproducing ex vivo and in vivo experiments^20,21^. Due to the importance of electrotonic interactions on the generation of phenomena such as alternans^22,23^ and afterdepolarisations^17,24^, it may be critically important in disease models to accurately capture the underlying coupling substrate, rather than implement a simple global reduction to cellular coupling.

Alternatively, other studies have modelled fibrosis by setting selected nodes to be unexcitable^25^, which introduces conduction heterogeneity more comparable to that presented in this study. Our approach enables conduction barriers to be captured without the need to remove nodes of tissue from being excitable. This, importantly, can promote temporary unidirectional conduction block which is a key mechanistic pathway for the generation of re-entrant circuits^26,27^. It is worth noting that many studies have further included electrotonic coupling between fibroblasts and myocytes in their descriptions of fibrosis^18,28,29^. This was not considered in the present study but is independent from the discretisation method and thus trivial to include, if desired, in future studies.

### 3.4 Comparison to other non-FDM or FEM approaches

Recent studies have presented alternatives to the established approaches for discretising cardiac tissue and solving the propagation of electrical excitation, many focusing on small tissue strands/slices at cellular-scale resolutions. Most directly comparable to our approach, network models of electrical coupling have been previously presented^25,30,31^. However, these models were not designed to implement complex anisotropy and are not directly set up for the modification of individual cellular (or nodal) connections, as in our approach.

At the cellular-scale, multiple recent studies have developed sophisticated models that describe intercellular electrical coupling to varying levels of detail. Some have explicitly described the time- and voltage-dependence of the gap junctional conductance, demonstrating these dynamic properties to be an important regulator of electrical conduction and repolarisation^13,16^. Jæger et al.^10^ presented a sophisticated model of intercellular coupling at the individual myocyte scale that accounts for the three domains of the extra-cellular, intra-cellular and membrane spaces. Such an approach enabled simulation of “micro-re-entry”, which cannot be reproduced using larger scale discretisations. Other studies have developed approaches that account for ephaptic coupling, in which ion channels localised to the intercalated discs play an important mechanistic role in electrical coupling^11,12^. These studies have demonstrated important dynamical implications for the features that were included, highlighting the disparity between cellular-scale and larger-scale tissue simulations. Translation of these models to the whole-heart scale is non-trivial, and there are many features that would need to be carefully considered (e.g., the relationship between coupling strength and discretisation spatial step; see “Limitations and further work” below). Nevertheless, compared to FDM and FEM, our new approach presents much greater compatibility with these more complex, dynamically regulated models: coupling is solved on the connection, not the node, and the gap-junction conductance of each connection is an explicit parameter which can be easily modified through the inclusion of gating variables, and extended to account for the conductivities in the different cellular/extracellular domains.

Very few studies have attempted to include these finer details of electrical coupling at the whole-heart scale. In Hurtado et al., (2020)^32^, non-linear homogenization theory was used to develop an up-scalable approximation of gap junctions that does not require explicit description of the underlying dynamics; in Saliani et al., (2022)^33^, a non-ohmic model of intercellular coupling was developed that was applied to a cable-based geometrical model, and included interruptions to intercellular coupling similar to our study. Both approaches offer powerful and complex alternatives to the established models of cardiac intercellular coupling. Our model represents a different philosophy based on simplicity and broad applicability; the ability to easily implement the model on structured grid geometries was a major motivation. It would be valuable in future studies to compare these different models and their resulting dynamics across a range of physiological and pathophysiological conditions.

### 3.5 Limitations and further work

Whereas this method is based on “intercellular connections”, the connection between nodes in a discretised geometry does not directly correspond to individual myocyte connections, as each node/voxel occupies a volume that contains multiple myocytes. This limitation is not unique to our new approach and applies to any tissue implementation that is not at the direct cellular scale. However, this does imply that modifying “intercellular connections” at this scale is not directly translatable to heterogeneous connections at the individual myocyte level. Nevertheless, it still enables implementation of functional barriers to conduction without requiring excitable nodes to be removed, which we argue is a more accurate representation of the underlying substrate. To further bridge this gap, future studies could develop high-throughput image analysis approaches to derive this model from local, high-resolution histological data (e.g., describing connexin expression and fibrosis) further to the whole-heart data for which it has been designed, and appropriate approximations could be developed in simulations to match models at different scales.

A further limitation, albeit another not specific to this model but rather of most approaches to modelling spatial coupling, is that the absolute coupling strength between two nodes is dependent on the spatial resolution such that a set conduction velocity can be matched. However, electrotonic coupling between myocytes in *reality* is clearly not dependent on the spatial resolution of a *geometrical model,* and this may have important implications on dynamics: electrotonic coupling affects cellular action potential morphology, including the rapid upstroke and the potential emergence of after depolarisations. Potential solutions to this limitation will be considered in future works.

Finally, there is a choice of how to treat mixed connections in the model (where one node contributes an axial but the other a transverse connection), and it is not clear which approach would be the most accurate. For example, a mixed connection would be observed at an abrupt change in fibre orientation, where the axial conductance of one node is coupled to the transverse of its neighbour. As presented in this study, the junctional conductance was simply an average of the two contributing conductances, independent of which type of connection they were. One could just as easily make a choice to model such junctions as either fully axial (axial connection only contributes to the junctional conductance) or fully transverse. This is a choice which could be carefully explored in future studies.

### 3.6 Conclusion

We have presented a simple alternative formulation to the reaction-diffusion equation which sums over gap junctional currents rather than discretising the continuous gradients. Whereas the model shares superficial similarities with FDM implementations, it also contains a fundamentally different underlying approach which presents advantages. Most notable is the ability to directly control intercellular connections, and suitability for integration with more sophisticated models of intercellular coupling mechanisms. These features enable more accurate and robust simulations of cardiac tissue to be performed, opening new avenues for the elucidation of underlying arrhythmia mechanisms.

## 4 Acknowledgements

The authors would like to thank Dr. Dominic Whittaker for helpful conversations during the early development of the approach, and Dr. Enric Alvarez Lacalle and Prof. Arun Holden for providing useful feedback on early versions of the manuscript.

## 7 Methods

### 7.1 Overall approach

Cardiac myocytes are electrically coupled through intercalated discs which are connections between cells that contain gap junctions enabling intercellular flow of ionic currents^34^. Myocytes are elongated in structure (with a length approximately five-to-ten times their diameter) and gap junctions are preferentially located at the longitudinal (or axial) connections, forming fibre-like strands of coupled myocytes. Coupling is also observed in the direction perpendicular to myocyte orientation but is less than in the axial direction, and the conduction velocity in the axial direction is typically at least twice that in the transverse direction^2^. This feature of differentiated axial and transverse intercellular coupling – and the ability to interrupt them individually - underlies the philosophy our proposed novel approach.

The approach operates on a structured gird of nodes i.e., a square (2D) or cuboid (3D) lattice. This represents the simplest discrete reconstruction of cardiac images, as opposed to an unstructured gird comprised of polygonal elements and vertices, and corresponds to the same system on which the FDM approach is applied. In this system, we present the inherently discrete equation governing the voltage differential for some node, *i*

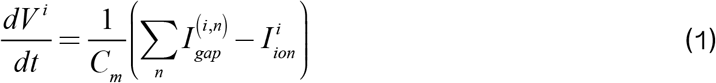

Where *I*_gap_^(i,n)^ is the current associated with gap junction *n* that is connected to node *i* and the sum over *n* corresponds to all junctions associated with node *i* (in 2D this is a sum of up to eight junctions; in 3D it is a sum of up to 26 junctions). It is common to absorb the membrane capacitance, *C*_m_, into the formulation of the ionic currents, through the definition of the maximal conductances in units of nS/pF. The same will be applied here for the conductance of the gap junctions, and so equation (1) simplifies to:

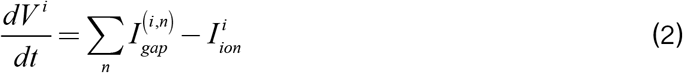

The junctional current for some junction, *n*, between two adjacent nodes, *i* and *j* is defined by:

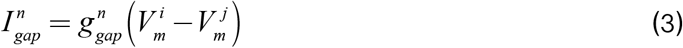

Where *g*_gap_^n^ is the conductance of gap junction *n*. The gap junctional current for the nodes *i* and *j* the term that contributes to the sum in equations (1) and (2), is therefore given by:

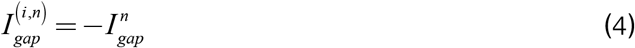

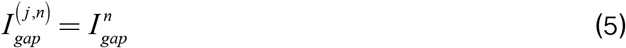

Note that we use the terms “gap junction”, “junctional current” and “junctional conductance” broadly here, as they may not correspond to individual gap junctions between individual *myocytes,* but rather the connections between *nodes.* Moreover, this approach is not a formulation of gap junctional dynamics itself, and the conductance of the junction will be assumed to be constant (i.e., not voltage- or time-dependent). Rather, we present a method to discretise the tissue model in order to determine the magnitude of this gap junctional conductance between neighbouring nodes based on local myocyte orientation.

We must now determine how nodes are connected to form junctions, and how to derive the gap junctional conductance, *g*_gap_^n^ based on local myocyte orientation. Throughout the remainder of the methods section for brevity, only the minimum number of equations necessary to understand the construction of the model will be shown, exploiting symmetries (e.g., x-directions will be shown but not the symmetric y and z equivalents); full equations are provided in the Online Supplement.

### 7.2 Derivation of the approach in 2D

#### 7.2.1 Axial and transverse connections

In a 2D structured grid, each node can be connected to eight other nodes, corresponding to four along the axes (±x, ±y) and four along the diagonals (±xy^++^, ±xy^+-^) (**Fig. 1a**). Due to the different directions along each of these lines being indifferent (+x is the same as -x), this reduces to four independent connection directions: x, y, xy^++^ and xy^+-^. Note that xy^++^ refers to the diagonal where the sign on x and y is identical (+x and +y, or -x and -y) and xy^+-^ to the diagonal where the signs are different.

The main concept of this approach is that, for each node, the contribution to the connection to its neighbours in each of these directions will be determined from weighting terms defined by the local myocyte orientation. Defining *g*_a_ as the conductance of a gap junction in the axial direction (along the primary myocyte orientation) and *g*_t_ as the conductance of a gap junction in the transverse direction (i.e., orthogonal to the primary myocyte orientation), the nodal junctional conductance in each direction (*g*_ei_^node^) can be described as:

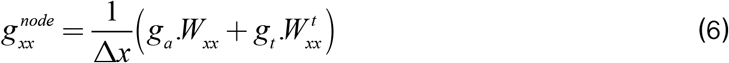

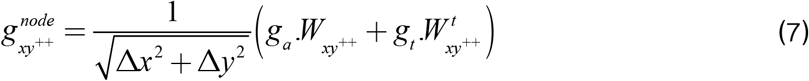

And similarly for the y and xy^+-^ directions (Online Supplement), where *g*_xx_^node^ is the contribution of the node to the gap junctional conductances in the x-direction, and equivalently for all other directions, *W*_xx_ is the weight towards the x-axis for the axial component, *W*_xx_^t^ is the weight towards the x-axis for the transverse component (and equivalent for all other directions), and Δx and Δy refer to the discretisation space step in each dimension and are included from geometrical arguments. Due to considerations regarding the relationship between coupling strength and the spatial step (see Discussion), it may also be desirable to define these conductances independent of the spatial steps. In this case, it is important to retain the factor of 1/√2 in the diagonal terms, or otherwise account for the increased distance in the diagonal directions, for geometrical consistency.

#### 7.2.2 Defining the junction conductance and currents

Before describing how the weighting terms are derived, it is important to relate the nodal conductances (equations 6–7) to the junctional conductance in equation (3). A junction is formed between all pairs of nodes which correspond to tissue, adjacent to each-other in any of the coupling directions. For some junction, *n*, the conductance, *g*_gap_^n^ in equation (3), is given by the mean of the nodal conductances in the direction in which they are coupled (**Fig. 1b**). For example, for two nodes, *i* and *j*, adjacent to each-other in the x-direction, forming a junction *n*

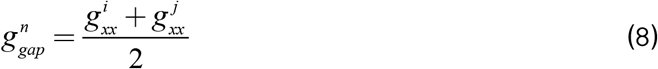

And equivalently for all other directions in which nodes may be coupled. There is no distinction here whether the connection is axial or transverse, so mixed connections (e.g., at an abrupt change in fibre) are possible. Every time a junction is created, maps must be created (for implementation purposes) defining which node is the positive and which is the negative contributor to this junction

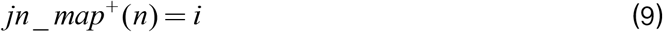

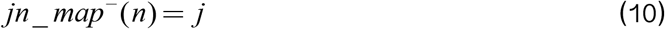

The gap junctional current for junction *n*, equation (3), is therefore:

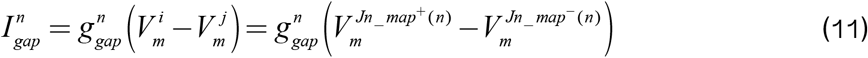

And these junctional currents are summed for each node as described in equations (2, 4 and 5).

#### 7.2.3 Deriving the weights based on myocyte orientation

The weights that scale the contribution of the axial and transverse gap junctional conductances to each nodal directional conductance term (equations 6–7) must be defined. The myocyte orientation in 2D will always point in a segment between one axis and one diagonal direction (**Extended Data Fig. 1**). Due to periodicities and symmetries in the trigonometric functions, we can consider and calculate the weight towards the axis and diagonal independent of in which segment the orientation is pointing. We can first calculate the angle from the x-axis, θ:

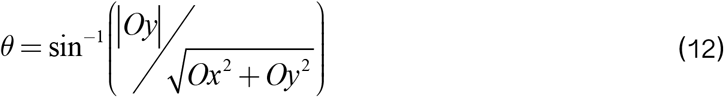

where *Ox* is the x-component of the normalised orientation vector, and *Oy* is the y-component. This will always return an angle between 0 rad (when the orientation is along the x axis) and π/2 rad (when the orientation is along the y axis). We can then define a weight which linearly depends on this angle as a measure of where the orientation lies between the x or y axis (θ = 0 or π/2 rad, respectively) and the diagonal (θ = π/4 rad):

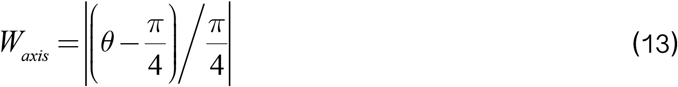

i.e., *W*_axis_ is equal to 1 if the orientation points exactly along either the x or y axis (θ = 0 or π/2 rad, respectively), equal to 0 if it points exactly along the diagonal (θ = π/4 rad), and linearly varies between 0 and 1 based on the angle in-between (**Extended Data Fig. 1**). The weight towards the diagonal is then simply:

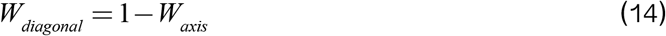

A conditional algorithm could then be applied to determine to which axis and diagonal the weighting terms should be applied. Alternatively, and perhaps more concisely, these choices can be absorbed into a single general equation through the introduction of binary pointing parameters. Thus, we may define:

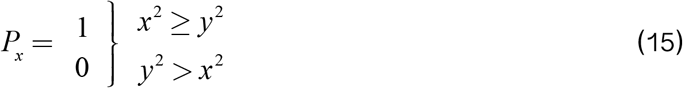

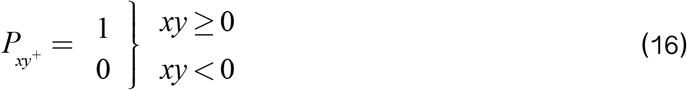

where *P*_x_ is a binary parameter which is equal to 1 if the orientation vector points between the x-axis and either diagonal and equals zero otherwise; *P*_xy+_ is equal to 1 if the orientation points between the diagonal xy^++^ and either axis. Note that *P*_y_ and *P*_xy-_ could also be introduced, defined as 1-P_x_ and 1-P_xy+_, respectively. From this, the weighting towards the four directions is then given by:

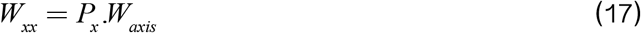

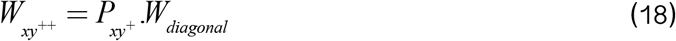

This approach can be illustrated by considering some simple example cases (Fig 1D).

#### 7.2.4 Transverse direction

A major feature of this approach is to apply the transverse coupling always at an orthogonal direction to the myocyte orientation (**Extended Data Fig. 1**), as opposed to FDM, for example, where the basic isotropic coupling is always applied along the axes, independent of the myocyte orientation. In 2D the transverse weights are trivial to calculate:

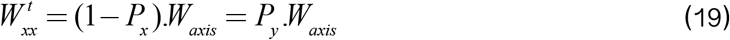

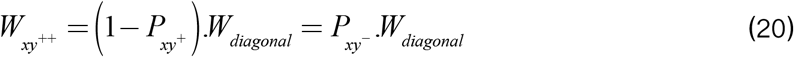

Which are the same as equations (17–19) with the pointing parameters swapped.

### 7.3 Derivation of the approach in 3D

#### 7.3.1 Primary myocyte orientation weights in 3D

In 3D each node can be connected to 26 neighbours (i.e., 13 independent directions given the +/- directional indifference), corresponding to the axes and diagonals in each plane and the four corners (**Fig. 1a**). The primary myocyte orientation vector therefore points in a segment which corresponds to a quadrant on the surface of a cube, i.e., contributing towards the weight of up to four directions at once (**Fig. 1c**). Similar to the approach in 2D, we can consider how the weights towards these four directions depend on the angle(s), independent of which axes and diagonals define the quadrant. The four directions (**Fig. 1c**) correspond to the axis, the in-plane diagonal (dependent on θ_1_), the elevation diagonal (dependent on θ_2_) and the corner (dependent on both θ_1_ and θ_2_). Note that both θ_1_ and θ_2_ are defined as the projection in the axis planes i.e., θ_2_ does not correspond to the spherical polar coordinate’s elevation angle. We can then define the weight towards each of these directions as (**Fig. 1c**):

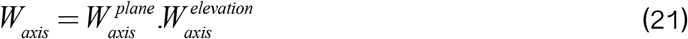

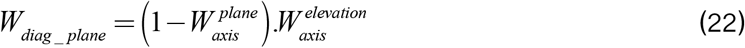

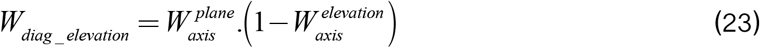

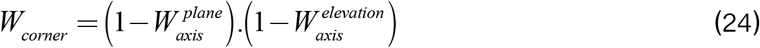

where *W*_axis_^plane^ and *W*_axis_^elevation^ are defined from the two angles, θ_1_ and θ_2_, in the same way as the 2D model (dependent on which axes contribute to these calculations). The three angles (θ_xy_, θ_xz_, θ_yz_) and corresponding weights (*W*_axis_^xy^, *W*_acis_^xz^, *W*_axis_^yz^) in each plane are defined by:

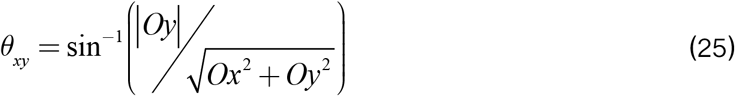

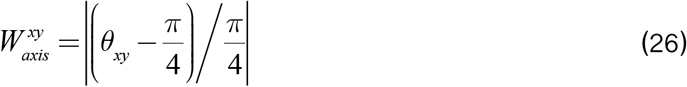

and the pointing parameters in 3D become:

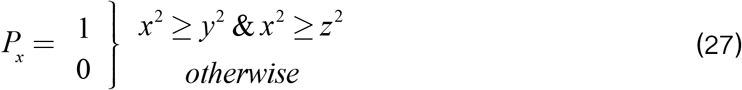

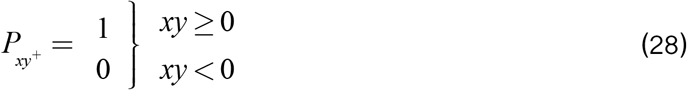

And similarly for the y- and z-directions (**Online Supplement**). Note that only one of *P*_x_, *P*_y_ and *P*_z_ can be non-zero at any time (the orientation can only be pointing primarily towards one of these axes). *P*_xy+_, *P*_xz+_ and *P*_yz+_ can all be non-zero. In full, therefore, the weights towards the four quadrant directions are given by:

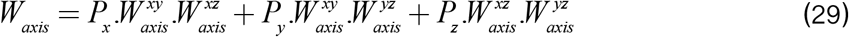

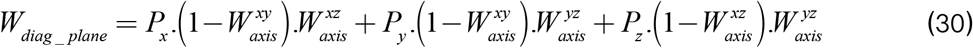

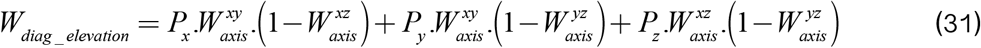

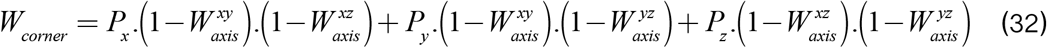

Note that because only one of Px, Py and Pz can be non-zero at a time, these equations reduce immediately to the equivalent of equations (23–26).

The weighting towards each direction is defined by:

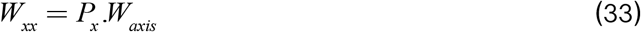

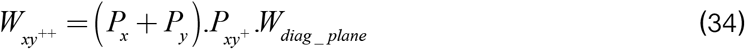

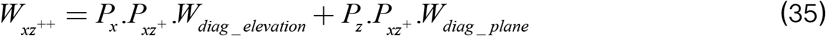

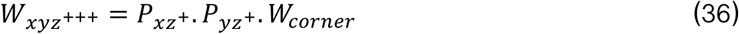

And similarly for all other directions (**Online Supplement**). Note that the form of equation (36) results from the convention that when pointing primarily towards z (*P*_z_ =1), the xz diagonal is considered “in plane” and the yz diagonal considered the “elevation” direction, whereas when pointing towards x or y, the xz and yz are both considered the elevation direction. This is an arbitrary choice, and the results would be the same whichever choice is made. As with the 2D case, when the pointing parameters are factored into these equations, only a maximum of four terms will result in being non-zero: one axis, two diagonals, and one corner.

#### 7.3.2 Transverse connections in 3D

In 3D, two transverse directions are necessary. These may correspond to the distinguishable “sheet” and “sheet normal” directions^35^ if these data describing the laminar/sheet structure of cardiac tissue are available [e.g. from diffusion tensor MRI^36–40^] or to two indistinguishable (equal magnitude) transverse directions if only myocyte orientation data are available (i.e. in the absence of data describing the laminar/sheet architecture of cardiac tissue). The junctional conductance is therefore given by:

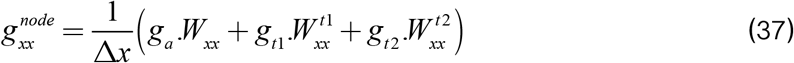

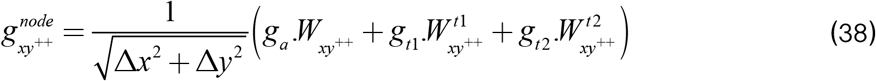

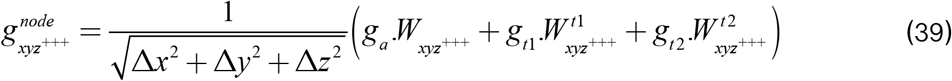

And similarly for all other directions (**Online Supplement**). Similarly to the 2D case, if these are defined independent of the spatial step, the factors of 1/√2 and 1/√3 must be retained for the diagonal and corner directions, respectively. If differentiating between in-sheet and transverse to sheet, then g_t1_ > g_t2_, whereas if not imposing this distinction, g_t1_ = g_t2_ = g_t_ and the equation(s) reduce to:

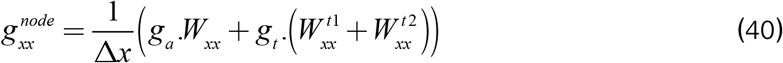

If implementation is based on three eigenvectors (e.g, from DTI), then the orientation components of the two transverse vectors directly determine *W*_ei_^11^ and *W*_ei_^12^ for each direction through the same equations (34–37) as the primary orientation vector (**Extended Data Fig. 1**). However, if only information on the primary eigenvector is defined, then we must define the transverse vectors from this (**Online Supplement**).

### 7.4 Implementing heterogeneous media

One main advantage, and indeed motivation for the development, of the above approach is to be able to modify cellular connections directly. For example, connexins (gap junctions) are heterogeneously expressed in tissue and may not exist between every adjacent myocyte. In disease in particular, highly heterogeneous media may be observed, corresponding to heterogeneity in connexin expression, or increased fibrosis. The latter may be associated with a loss of cellular connectivity and changes to the anisotropy ratio of conduction velocity, implying differential loss of transverse and axial connections.

Implementing these heterogeneities in this network model may be approached in different ways, depending on the application. For example, a map can be passed in that simply scales local (nodal) ga and gt, in the same way as one might do when using FDM. However, we can also scale the *junctions* directly, or even remove them. In this study we demonstrate these different approaches and their potential impact on function, although it should be clarified that we are not constructing physiologically validated parameter combinations for different health and disease states but, rather, demonstrating how this model may be applied in these contexts.

This approach readily facilitates differential removal of axial and transverse cellular connections. One can define a threshold for each type of connection that determines the probability of a connection being removed. It is important to identify if a connection is axial (the contribution from both nodes is from the primary orientation vector), transverse (the contribution from both nodes is from a transverse vector) or mixed (one node contributes an axial component and the other a transverse). Once complete, random numbers can be generated and combined with the threshold to determine which connections are removed. Alternatively, as a more sophisticated approach, junctional conductance could be scaled by continuous factors, for example, randomly sampled from various user-defined distributions or based on analysis of variability in experiment.

### 7.5 Cellular dynamics and model parameters

Example simulations presented in this study were performed using cellular electrophysiology described by the hybrid minimal model presented in Colman (2019). This is a simple model parameterised for the magnitudes of ionic currents and voltage range observed in cardiac electrical excitation. It is useful and desirable to be able to define the connection parameter conductances (*g*_a_ and *g*_t_) from the diffusion coefficient, *D*, to enable easy implementation to replace FDM, or similar, where *D* has been defined to match conduction velocity. A conversion factor can be obtained through analysis of a simple case (**Online Supplement**), yielding:

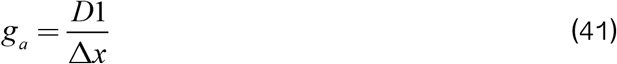

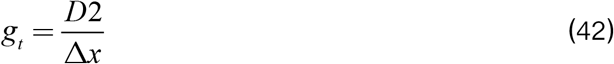

As will be shown in results, this conversion gives the same conduction velocities as the FDM method in these simple conditions. Note that this linear relationship means *g*_t_ can be defined from the anisotropy ratio (AR) and *g*_a_, in the same way as often for D1 and D2:

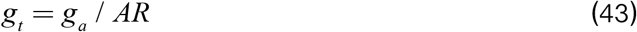

In this setup, the units of *g*_a_ and *g*_t_ are the same as those for the maximal conductances of the ion currents (nS/pF), as the membrane capacitance (*C*_m_) in equation (1) has been absorbed into these maximal conductance terms (equation 2). However, this model does not require this to be the case and enables coupling between cells of different *C*_m_ while still conserving current; one could define *g*_a_ as an absolute conductance (nS) and maintain inclusion of the factor *C*_m_^-1^ when implementing equation (1).

**Extended Data Fig 1.**
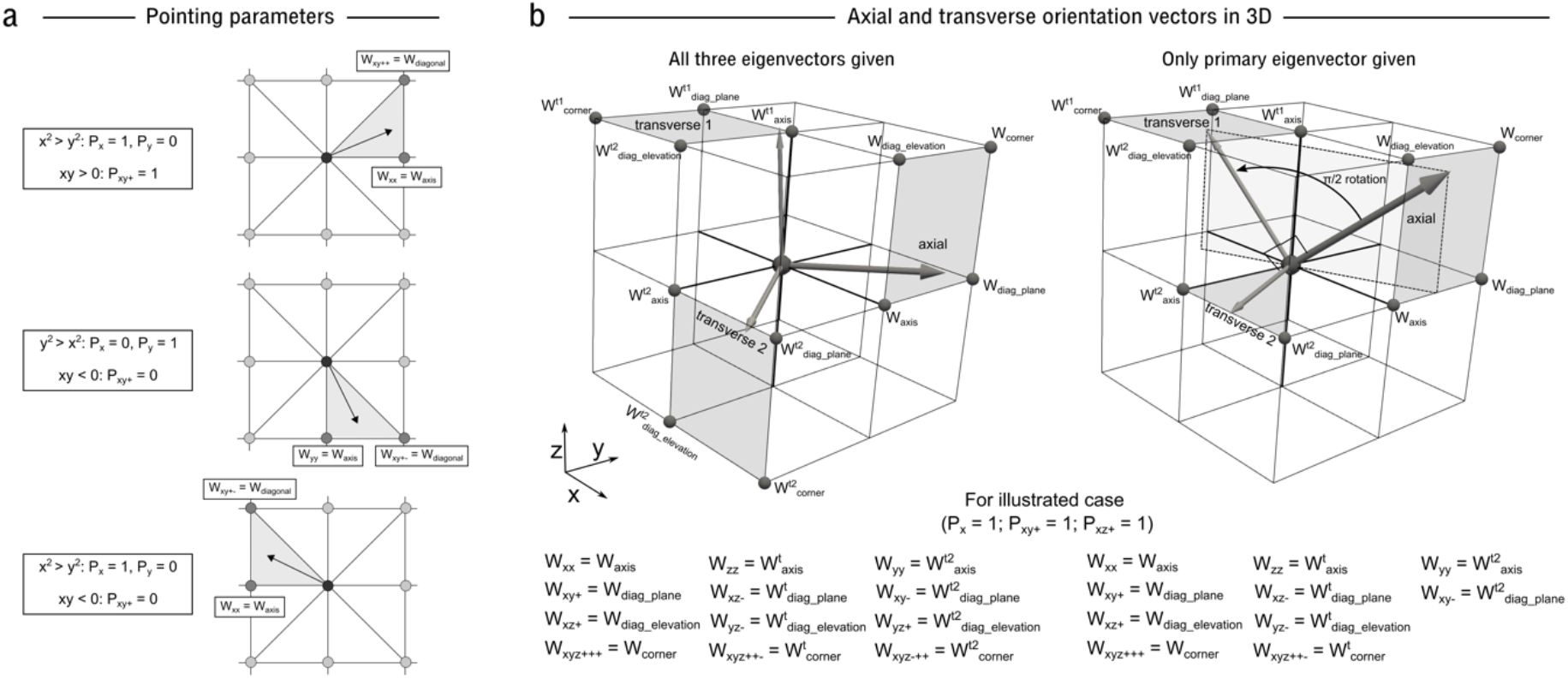
Illustration of pointing parameters and transverse connections. **a** – illustration of the different values the binary pointing parameters take dependent on in which segment the myocyte orientation points. **b** – illustration of the transverse orientation vectors and the directions to which they contribute in 3D. **Left** - illustration of the three orientation vectors as given by imaging data; each vector points in a quadrant, contributing to up to four directions. **Right** – illustration of the definition of the two transverse vectors if only the primary eigenvector is given. From the axial orientation, a π/2 rotation is applied towards the z-axis to define transverse vector 1. The cross-product of these two vectors then defines transverse vector 2, which will always point in the x-y plane.

