## Supplementary material for "A simple approach for image-based modelling of the heart that enables robust simulation of highly heterogeneous electrical excitation": Online Supplement

*Model Derivation, Verification, and Symmetry Considerations*

Michael A Colman & Alan P Benson

School of Biomedical Sciences, Faculty of Biological Sciences, University of Leeds, UK

### 1 Methods

#### 1.1 Overall approach

For the purposes of clarity and convenience, the full model derivation will be given here, including repetition of information already provided in the manuscript.

The approach operates on a structured gird of nodes i.e., a square (2D) or cuboid (3D) lattice. This represents the simplest discrete reconstruction of cardiac images, as opposed to an unstructured gird comprised of polygonal elements and vertices, and corresponds to the same system on which the FDM approach is applied. In this system, we present the inherently discrete equation for some node, *i*, corresponding to excitable tissue within the structured spatial grid

$$\frac{dV^i}{dt} = \frac{1}{C_m}\left(\sum_n I_{gap}^{(i,n)} - I_{ion}^i\right) \quad (1)$$

Where $I_{gap}^{(i,n)}$ is the current associated with gap junction *n* that is connected to node *i*, and the sum over *n* corresponds to all junctions associated with node *i* (in 2D this is a sum of up to eight junctions; in 3D it is a sum of up to 26 junctions). It is common to absorb the membrane capacitance, $C_m$, into the formulation of the ionic currents, through the definition of the maximal conductances in units of nS/pF. The same will be applied here for the conductance of the gap junctions, and so equation (1) simplifies to:

$$\frac{dV^i}{dt} = \sum_n I_{gap}^{(i,n)} - I_{ion}^i \quad (2)$$

The junctional current for some junction, *n*, between two adjacent nodes, *i* and *j*, is defined by:

$$I_{gap}^n = g_{gap}^n\left(V_m^i - V_m^j\right) \quad (3)$$

Where $g_{gap}^n$ is the conductance of gap junction *n*. The gap junctional current for the nodes *i* and *j*, the term that contributes to the sum in equations (1) and (2), is therefore given by:

$$I_{gap}^{(i,n)} = -I_{gap}^{n} \quad (4)$$

$$I_{gap}^{(j,n)} = I_{gap}^{n} \quad (5)$$

Note that we use the terms “gap junction”, “junctional current” and “junctional conductance” broadly here, as they do not correspond to individual gap junctions between individual myocytes, but rather the connections between nodes. Moreover, this approach is not a formulation of gap junctional dynamics itself, and the conductance of the junction will be assumed to be constant (i.e., not voltage or time dependent). Rather, we present a method to discretise the tissue model based on local myocyte orientation in order to determine the magnitude of this gap junctional conductance between neighbouring nodes.

We must now determine how nodes are connected to form junctions, and how to derive the gap junctional conductance, $g_{gap}^{n}$ based on local myocyte orientation.

#### 1.2 Derivation of the approach in 2D

##### 1.2.1 Axial and transverse connections

In a 2D structured grid, each node can be connected to eight other nodes, corresponding to four along the axes (±x, ±y) and four along the diagonals ($\pm xy^{++}$, $\pm xy^{+-}$) (**Fig. S1A**). Due to the different directions along each of these lines being indifferent (+x is the same as -x), this reduces to four independent connection directions: x, y, $xy^{++}$ and $xy^{+-}$. Note that $xy^{++}$ refers to the diagonal where the sign on x and y is identical (+x and +y, or -x and -y) and $xy^{+-}$ to the diagonal where the signs are different.

The main concept of this approach is that, for each node, the contribution to the connection to its neighbours in each of these directions will be determined from weighting terms defined by the local myocyte orientation. Defining $g_a$ as the conductance of a gap junction in the axial direction (along the primary myocyte orientation) and $g_t$ as the conductance of a gap junction in the transverse direction (i.e., orthogonal to the primary myocyte orientation), the nodal junctional conductance in each direction ($g_{ei}^{node}$) can be described by:

$$g_{xx}^{node} = \frac{1}{\Delta x}\left(g_a.W_{xx} + g_t.W_{xx}^{t}\right) \quad (6)$$

$$g_{yy}^{node} = \frac{1}{\Delta y}\left(g_a.W_{yy} + g_t.W_{yy}^{t}\right) \quad (7)$$

$$g_{xy^{++}}^{node} = \frac{1}{\sqrt{\Delta x^2 + \Delta y^2}}\left(g_a.W_{xy^{++}} + g_t.W_{xy^{++}}^{t}\right) \quad (8)$$

$$g_{xy^{+-}}^{node} = \frac{1}{\sqrt{\Delta x^2 + \Delta y^2}}\left(g_a.W_{xy^{+-}} + g_t.W_{xy^{+-}}^{t}\right) \quad (9)$$

where $g_{xx}^{node}$ is the contribution of the node to the gap junctional conductances in the x-direction, and equivalently for all other directions, $W_{xx}$ is the weight towards the x-axis for the axial component, $W_{xx}^{t}$ is the weight towards the x-axis for the transverse component (and equivalent for all other directions), and Δx and Δy refer to the discretisation space step in each dimension and are included from geometrical arguments. Due to considerations regarding the relationship between coupling strength and the spatial step (see Discussion), it may also be desirable to define these conductances independent of the spatial steps. In this case, it is important to retain the factor of 1/√2 in the diagonal terms, or otherwise account for the increased distance in the diagonal directions, for geometrical consistency.

##### 1.2.2 Defining the junction conductance and currents

Before describing how the weighting terms are derived, it is important to relate the nodal conductances (equations 6-9) to the junctional conductance and currents, which contribute to the sum of equations (1-2). A junction is formed between all pairs of nodes which correspond to tissue, adjacent to each-other in any of the coupling directions. For some junction, *n*, the conductance, $g_{gap}^{n}$ in equation (3), is given by the mean of the nodal conductances in the direction in which they are coupled (**Fig. S1B**). For example, for two nodes, *i* and *j*, adjacent to each-other in the x-direction, forming a junction *n*:

$$g_{gap}^{n} = \frac{g_{xx}^{i} + g_{xx}^{j}}{2} \tag{10}$$

And equivalently for all other directions in which nodes may be coupled. There is no distinction here whether the connection is axial or transverse, so mixed connections (e.g., at an abrupt change in fibre) are perfectly possible. Every time a junction is created, maps must be created (for implementation purposes) defining which node is the positive and which is the negative contributor to this junction:

$$jn\_map^{+}(n) = i \tag{11}$$

$$jn\_map^{-}(n) = j \tag{12}$$

The gap junctional current for junction *n*, equation (3), is therefore:

$$I_{gap}^{n} = g_{gap}^{n}\left(V_{m}^{i} - V_{m}^{j}\right) = g_{gap}^{n}\left(V_{m}^{Jn\_map^{+}(n)} - V_{m}^{Jn\_map^{-}(n)}\right) \tag{13}$$

And these junctional currents are summed for each node as described in equations (2, 4 and 5).

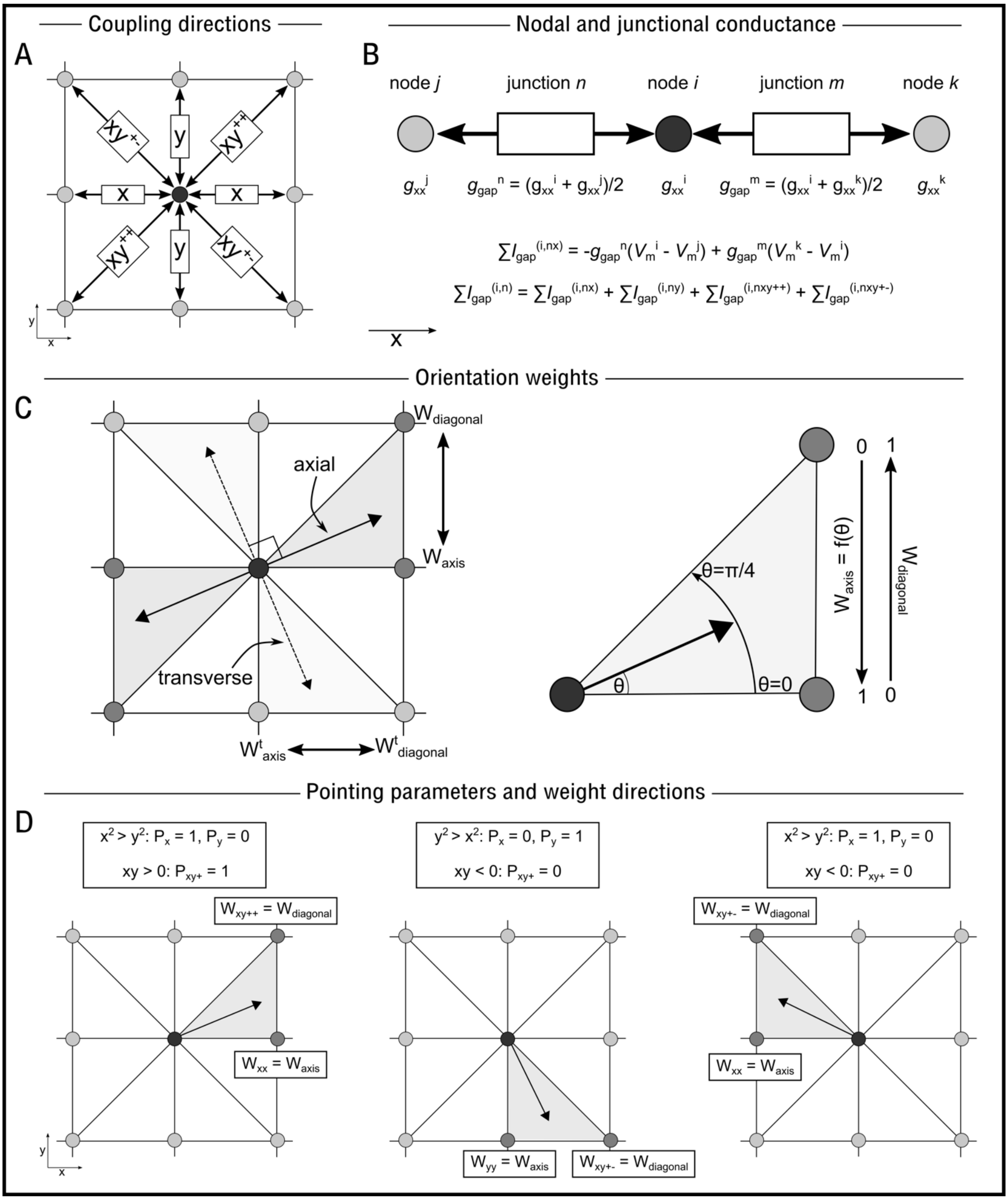


**Figure S1.** Illustration of the model connections in 2D. **A** – illustration of the different coupling directions and terminology used in this paper. **B** – illustration of how the nodal and junctional conductance parameters are related and contribute to the total gap junctional current, for three nodes *i*, *j*, and *k* and two junctions *n* and *m*. **C** – illustration of the relationship between myocyte orientation angle and the weighting terms towards the axis and diagonal; in this particular example, the myocyte orientation is between the positive x axis and the $xy^{++}$ diagonal. **D** – illustration of the different values the binary pointing parameters take dependent on in which segment the myocyte orientation points.

##### 1.2.3 Deriving the weights based on myocyte orientation

Now we must determine the weights that scale the contribution of the axial and transverse gap junctional conductances to each nodal directional conductance term (equations 6-9). The myocyte orientation in 2D will always point in a segment between one axis and one diagonal direction (**Fig. S1C**). Due to periodicities and symmetries in the trigonometric functions, we can consider and calculate the weight towards the axis and diagonal independent of in which segment the orientation is pointing. We can first calculate the angle from the x axis, θ:

$$\theta = \sin^{-1}\left( |Oy| \Big/ \sqrt{Ox^2 + Oy^2} \right) \tag{14}$$

where $Ox$ is the x-component of the normalised orientation vector, and $Oy$ is the y-component. This will always return an angle between 0 rad (when the orientation is along the x axis) and π/2 rad (when the orientation is along the y axis). We can then define a weight which linearly depends on this angle as a measure of where the orientation lies between the x or y axis (θ = 0 or π/2 rad, respectively) and the diagonal (π/4 rad):

$$W_{axis} = \left| \left( \theta - \frac{\pi}{4} \right) \Big/ \frac{\pi}{4} \right| \tag{15}$$

i.e., $W_{axis}$ is equal to 1 if the orientation points exactly along either the x or y axis (θ = 0 or π/2 rad, respectively), equal to 0 if it points exactly along the diagonal (θ = π/4 rad), and linearly varies between 0 and 1 based on the angle in-between (**Fig. S1C**). The weight towards the diagonal is then simply:

$$W_{diagonal} = 1 - W_{axis} \tag{16}$$

A conditional algorithm could then be applied to determine to which axis and diagonal the weighting terms should be applied. Alternatively, and perhaps more concisely, these choices can be absorbed into a single general equation through the introduction of binary pointing parameters. Thus, we may define:

$$P_x = \begin{cases} 1 & x^2 \geq y^2 \\ 0 & y^2 > x^2 \end{cases} \tag{17}$$

$$P_{xy^+} = \begin{cases} 1 & xy \geq 0 \\ 0 & xy < 0 \end{cases} \tag{18}$$

where $P_x$ is a binary parameter which is equal to 1 if the orientation vector points between the x-axis and either diagonal and equals zero otherwise; $P_{xy+}$ is equal to 1 if the orientation points between the diagonal $xy^{++}$ and either axis. Note that $P_y$ and $P_{xy-}$ could also be introduced, defined as $1-P_x$ and $1-P_{xy+}$, respectively. From this, the weighting towards the four directions is then given by:

$$W_{xx} = P_x.W_{axis} \quad (19)$$

$$W_{yy} = P_y.W_{axis} = (1 - P_x).W_{axis} \quad (20)$$

$$W_{xy^{++}} = P_{xy^{+}}.W_{diagonal} \quad (21)$$

$$W_{xy^{+-}} = P_{xy^{-}}.W_{diagonal} = \left(1 - P_{xy^{+}}\right).W_{diagonal} \quad (22)$$

This approach can be illustrated by considering some simple example cases (**Fig. S1D**). If, for example, θ is between 0 and π/4 rad, then $x^2$ is larger than $y^2$ and thus $P_x = 1$ and $P_y = 0$; x and y are both positive and thus $xy > 0$ and $P_{xy+} = 1$. For this particular example, the weighting equations would reduce to:

$$W_{xx} = W_{axis} \quad (23)$$

$$W_{yy} = 0 \quad (24)$$

$$W_{xy^{++}} = W_{diagonal} \quad (25)$$

$$W_{xy^{+-}} = 0 \quad (26)$$

Further examples are illustrated in **Fig. S1D.**

##### 1.2.4 Transverse direction

A major feature of this approach is to apply the transverse coupling always at an orthogonal direction to the myocyte orientation (**Fig. S1C**), as opposed to FDM, for example, where the basic isotropic coupling is always applied along the axes, independent of the myocyte orientation. In 2D the transverse weights are trivial to calculate:

$$W^t_{xx} = (1 - P_x).W_{axis} = P_y.W_{axis} \quad (27)$$

$$W^t_{yy} = (1 - P_y).W_{axis} = P_x.W_{axis} \quad (28)$$

$$W_{xy^{++}} = \left(1 - P_{xy^{+}}\right).W_{diagonal} = P_{xy^{-}}.W_{diagonal} \quad (29)$$

$$W_{xy^{+-}} = \left(1 - P_{xy^{-}}\right).W_{diagonal} = P_{xy^{+}}.W_{diagonal} \quad (30)$$

Which are of course the same as equations (19-22) with the pointing parameters swapped.

#### 1.3 Derivation of the approach in 3D

##### 1.3.1 Primary myocyte orientation weights in 3D

In 3D each node can be connected to 26 neighbours (i.e., 13 independent directions given the +/- directional indifference), corresponding to the axes and diagonals in each plane and the four corners (**Fig. S2A**). The primary myocyte orientation vector therefore points in a segment which corresponds to a quadrant on the surface of a cube, i.e., contributing towards the weight of up to four directions at once (**Fig. S2B**). Similarly to the approach in 2D, we can consider how the weights towards these four directions depend on the angle(s), independent of which axes and diagonals define the quadrant. The four directions (**Fig. S2B**) correspond to the axis, the in-plane diagonal (dependent on $\theta_1$, defined in **Fig. S2B**), the elevation diagonal (dependent on $\theta_2$, again defined in **Fig. S2B**) and the corner (dependent on both $\theta_1$ and $\theta_2$). Note that both $\theta_1$ and $\theta_2$ are defined as the projection in the axis planes i.e., $\theta_2$ does not correspond to the spherical polar coordinate's elevation angle. We can then define the weight towards each of these directions as (**Fig. S2B**):

$$W_{axis} = W_{axis}^{plane} . W_{axis}^{elevation} \tag{31}$$

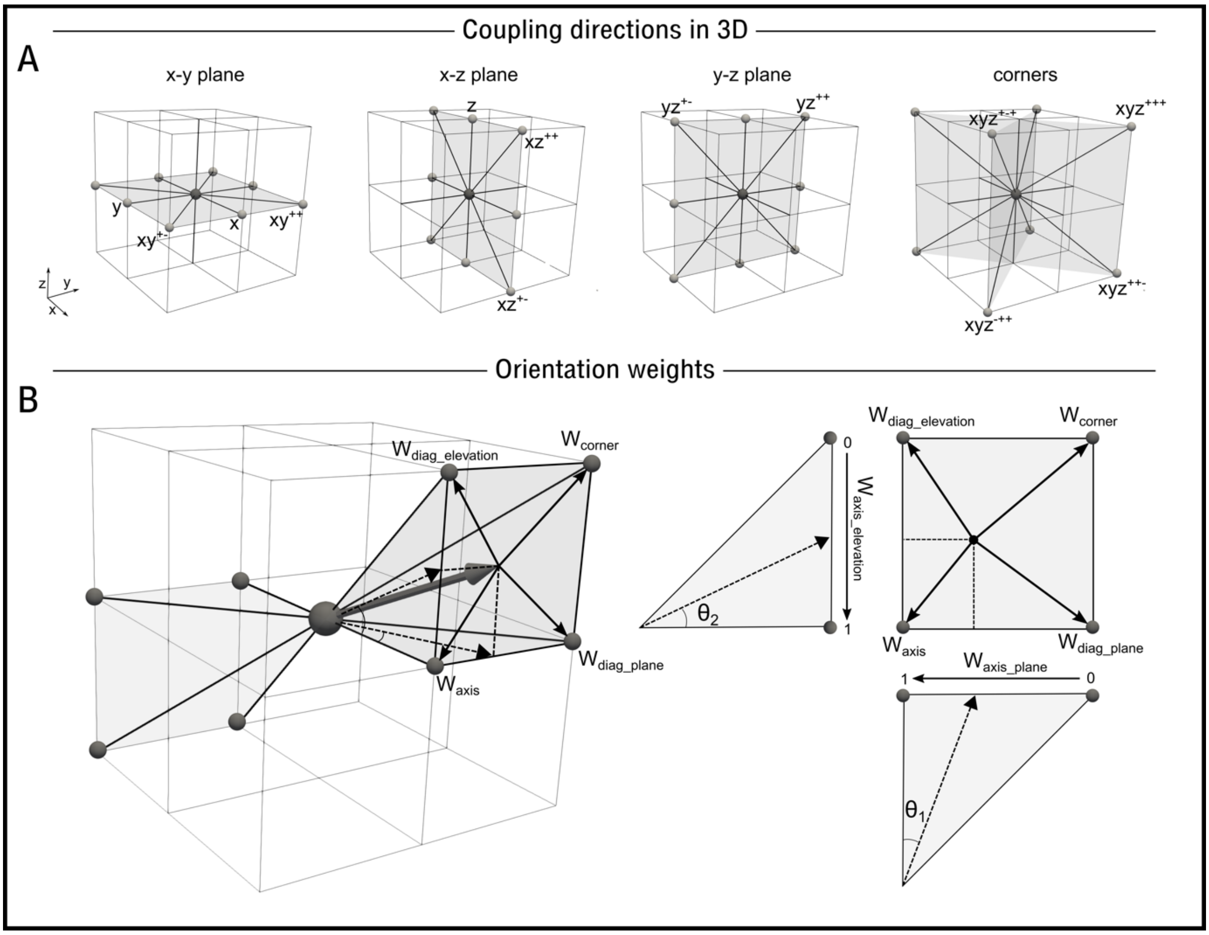


**Figure S2.** Illustration of coupling directions and orientation weights in 3D**. A** – The 13 different coupling directions (nodal connections) in a 3D structured grid. **B** – illustration of how the weight towards each direction in a quadrant is calculated.

$$W_{diag\_plane} = \left(1 - W_{axis}^{plane}\right) . W_{axis}^{elevation} \quad (32)$$

$$W_{diag\_elevation} = W_{axis}^{plane} . \left(1 - W_{axis}^{elevation}\right) \quad (33)$$

$$W_{corner} = \left(1 - W_{axis}^{plane}\right) . \left(1 - W_{axis}^{elevation}\right) \quad (34)$$

where $W_{axis}^{plane}$ and $W_{axis}^{elevation}$ are defined from the two angles, $\theta_1$ and $\theta_2$, in the same way as the 2D model (dependent on which axes contribute to these calculations). The three angles ($\theta_{xy}$, $\theta_{xz}$, $\theta_{yz}$) and corresponding weights ($W_{axis}^{xy}$, $W_{axis}^{xz}$, $W_{axis}^{yz}$) in each plane are defined by:

$$\theta_{xy} = \sin^{-1}\left( |Oy| \Big/ \sqrt{Ox^2 + Oy^2} \right) \quad (35)$$

$$\theta_{xz} = \sin^{-1}\left( |Oz| \Big/ \sqrt{Ox^2 + Oz^2} \right) \quad (36)$$

$$\theta_{yz} = \sin^{-1}\left( |Oz| \Big/ \sqrt{Oz^2 + Oy^2} \right) \quad (37)$$

$$W_{axis}^{xy} = \left| \left( \theta_{xy} - \frac{\pi}{4} \right) \Big/ \frac{\pi}{4} \right| \quad (38)$$

$$W_{axis}^{xz} = \left| \left( \theta_{xz} - \frac{\pi}{4} \right) \Big/ \frac{\pi}{4} \right| \quad (39)$$

$$W_{axis}^{yz} = \left| \left( \theta_{yz} - \frac{\pi}{4} \right) \Big/ \frac{\pi}{4} \right| \quad (40)$$

and the pointing parameters in 3D become:

$$P_x = \left. \begin{matrix} 1 \\ 0 \end{matrix} \right\} \begin{matrix} x^2 \geq y^2 \ \& \ x^2 \geq z^2 \\ otherwise \end{matrix} \quad (41)$$

$$P_y = \left. \begin{matrix} 1 \\ 0 \end{matrix} \right\} \begin{matrix} y^2 > x^2 \ \& \ y^2 \geq z^2 \\ otherwise \end{matrix} \quad (42)$$

$$P_z = \begin{matrix} 1 \\ 0 \end{matrix} \Bigg\} \begin{matrix} z^2 > x^2 \,\&\, z^2 > y^2 \\ otherwise \end{matrix} \quad (43)$$

$$P_{xy^+} = \begin{matrix} 1 \\ 0 \end{matrix} \Bigg\} \begin{matrix} xy \geq 0 \\ xy < 0 \end{matrix} \quad (44)$$

$$P_{xz^+} = \begin{matrix} 1 \\ 0 \end{matrix} \Bigg\} \begin{matrix} xz \geq 0 \\ xz < 0 \end{matrix} \quad (45)$$

$$P_{yz^+} = \begin{matrix} 1 \\ 0 \end{matrix} \Bigg\} \begin{matrix} yz \geq 0 \\ yz < 0 \end{matrix} \quad (46)$$

Note that only one of $P_x$, $P_y$ and $P_z$ can be non-zero at any time (the orientation can only be pointing primarily towards one of these axes). $P_{xy+}$, $P_{xz+}$ and $P_{yz+}$ can all be non-zero. In full, therefore, the weights towards the four quadrant directions are given by:

$$W_{axis} = P_x.W_{axis}^{xy}.W_{axis}^{xz} + P_y.W_{axis}^{xy}.W_{axis}^{yz} + P_z.W_{axis}^{xz}.W_{axis}^{yz} \quad (47)$$

$$W_{diag\_plane} = P_x.\left(1-W_{axis}^{xy}\right).W_{axis}^{xz} + P_y.\left(1-W_{axis}^{xy}\right).W_{axis}^{yz} + P_z.\left(1-W_{axis}^{xz}\right).W_{axis}^{yz} \quad (48)$$

$$W_{diag\_elevation} = P_x.W_{axis}^{xy}.\left(1-W_{axis}^{xz}\right) + P_y.W_{axis}^{xy}.\left(1-W_{axis}^{yz}\right) + P_z.W_{axis}^{xz}.\left(1-W_{axis}^{yz}\right) \quad (49)$$

$$W_{corner} = P_x.\left(1-W_{axis}^{xy}\right).\left(1-W_{axis}^{xz}\right) + P_y.\left(1-W_{axis}^{xy}\right).\left(1-W_{axis}^{yz}\right) + P_z.\left(1-W_{axis}^{xz}\right).\left(1-W_{axis}^{yz}\right) \quad (50)$$

Note that because only one of Px, Py and Pz can be non-zero at a time, these equations reduce immediately to the equivalent of equations (31-34).

The weighting towards each direction is defined by:

$$W_{xx} = P_x.W_{axis} \quad (51)$$

$$W_{yy} = P_y.W_{axis} \quad (52)$$

$$W_{zz} = P_z.W_{axis} \quad (53)$$

$$W_{xy^{++}} = \left(P_x + P_y\right).P_{xy^+}.W_{diag\_plane} \quad (54)$$

$$W_{xy^{+-}} = \left(P_x + P_y\right).\left(1 - P_{xy^+}\right).W_{diag\_plane} \quad (55)$$

$$W_{xz^{++}} = P_x . P_{xz^+} . W_{diag\_elevation} + P_z . P_{xz^+} . W_{diag\_plane} \tag{56}$$

$$W_{xz^{+-}} = P_x . \left(1 - P_{xz^+}\right) . W_{diag\_elevation} + P_z . \left(1 - P_{xz^+}\right) . W_{diag\_plane} \tag{57}$$

$$W_{yz^{++}} = \left(P_y + P_z\right) . P_{yz^+} . W_{diag\_elevation} \tag{58}$$

$$W_{yz^{+-}} = \left(P_y + P_z\right) . \left(1 - P_{yz^+}\right) . W_{diag\_elevation} \tag{59}$$

$$W_{xyz^{+++}} = P_{xz^+} . P_{yz^+} . W_{diag\_diag} \tag{60}$$

$$W_{xyz^{++-}} = P_{xy^+} . \left(1 - P_{xz^+}\right) . W_{diag\_diag} \tag{61}$$

$$W_{xyz^{+-+}} = \left(1 - P_{xy^+}\right) . P_{xz^+} . W_{diag\_diag} \tag{62}$$

$$W_{xyz^{-++}} = \left(1 - P_{xy^+}\right) . \left(1 - P_{xz^+}\right) . W_{diag\_diag} \tag{63}$$

Note that the form of equation (56-57) results from the convention that when pointing primarily towards z ($P_z$ =1), the xz diagonal is considered "in plane" and the yz diagonal considered the "elevation" direction, whereas when pointing towards x or y, the xz and yz are both considered the elevation direction. This is an arbitrary choice, and the results would be the same whichever choice is made. As with the 2D case, when the pointing parameters are factored into these equations, only a maximum of four terms will result in being non-zero: one axis, two diagonals, and one corner.

##### 1.3.2 Transverse connections in 3D

In 3D, two transverse directions are necessary. These may correspond to the distinguishable "sheet" and "sheet normal" directions (LeGrice et al., 1995) if these data describing the laminar/sheet structure of cardiac tissue are available [e.g. from diffusion tensor MRI (Helm et al., 2005; Holmes et al., 2000; Hsu et al., 1998; Scollan et al., 1998; Tseng et al., 2003)] or to two indistinguishable (equal magnitude) transverse directions if only myocyte orientation data are available (i.e. in the absence of data describing the laminar/sheet architecture of cardiac tissue). The junctional conductance is therefore given by:

$$g_{xx}^{node} = \frac{1}{\Delta x}\left(g_a . W_{xx} + g_{t1} . W_{xx}^{t1} + g_{t2} . W_{xx}^{t2}\right) \tag{64}$$

$$g_{yy}^{node} = \frac{1}{\Delta y}\left(g_a . W_{yy} + g_{t1} . W_{yy}^{t1} + g_{t2} . W_{yy}^{t2}\right) \tag{65}$$

$$g_{zz}^{node} = \frac{1}{\Delta z}\left(g_a . W_{zz} + g_{t1} . W_{zz}^{t1} + g_{t2} . W_{zz}^{t2}\right) \tag{66}$$

$$g^{node}_{xy^{++}} = \frac{1}{\sqrt{\Delta x^2 + \Delta y^2}}\left(g_a.W_{xy^{++}} + g_{t1}.W^{t1}_{xy^{++}} + g_{t2}.W^{t2}_{xy^{++}}\right) \tag{67}$$

$$g^{node}_{xy^{+-}} = \frac{1}{\sqrt{\Delta x^2 + \Delta y^2}}\left(g_a.W_{xy^{+-}} + g_{t1}.W^{t1}_{xy^{+-}} + g_{t2}.W^{t2}_{xy^{+-}}\right) \tag{68}$$

$$g^{node}_{xz^{++}} = \frac{1}{\sqrt{\Delta x^2 + \Delta z^2}}\left(g_a.W_{xz^{++}} + g_{t1}.W^{t1}_{xz^{++}} + g_{t2}.W^{t2}_{xz^{++}}\right) \tag{69}$$

$$g^{node}_{xz^{+-}} = \frac{1}{\sqrt{\Delta x^2 + \Delta z^2}}\left(g_a.W_{xz^{+-}} + g_{t1}.W^{t1}_{xz^{+-}} + g_{t2}.W^{t2}_{xz^{+-}}\right) \tag{70}$$

$$g^{node}_{yz^{++}} = \frac{1}{\sqrt{\Delta y^2 + \Delta z^2}}\left(g_a.W_{yz^{++}} + g_{t1}.W^{t1}_{yz^{++}} + g_{t2}.W^{t2}_{yz^{++}}\right) \tag{71}$$

$$g^{node}_{yz^{+-}} = \frac{1}{\sqrt{\Delta y^2 + \Delta z^2}}\left(g_a.W_{yz^{+-}} + g_{t1}.W^{t1}_{yz^{+-}} + g_{t2}.W^{t2}_{yz^{+-}}\right) \tag{72}$$

$$g^{node}_{xyz^{+++}} = \frac{1}{\sqrt{\Delta x^2 + \Delta y^2 + \Delta z^2}}\left(g_a.W_{xyz^{+++}} + g_{t1}.W^{t1}_{xyz^{+++}} + g_{t2}.W^{t2}_{xyz^{+++}}\right) \tag{73}$$

$$g^{node}_{xyz^{++-}} = \frac{1}{\sqrt{\Delta x^2 + \Delta y^2 + \Delta z^2}}\left(g_a.W_{xyz^{++-}} + g_{t1}.W^{t1}_{xyz^{++-}} + g_{t2}.W^{t2}_{xyz^{++-}}\right) \tag{74}$$

$$g^{node}_{xyz^{+-+}} = \frac{1}{\sqrt{\Delta x^2 + \Delta y^2 + \Delta z^2}}\left(g_a.W_{xyz^{+-+}} + g_{t1}.W^{t1}_{xyz^{+-+}} + g_{t2}.W^{t2}_{xyz^{+-+}}\right) \tag{75}$$

$$g^{node}_{xyz^{-++}} = \frac{1}{\sqrt{\Delta x^2 + \Delta y^2 + \Delta z^2}}\left(g_a.W_{xyz^{-++}} + g_{t1}.W^{t1}_{xyz^{-++}} + g_{t2}.W^{t2}_{xyz^{-++}}\right) \tag{76}$$

Similarly to the 2D case, if these are defined independent of the spatial step, the factors of 1/√2 and 1/√3 must be retained for the diagonal and corner directions, respectively. If differentiating between in-sheet and transverse to sheet, then $g_{t1} > g_{t2}$, whereas if not imposing this distinction, $g_{t1} = g_{t2} = g_t$ and the equation(s) reduce to:

$$g^{node}_{xx} = \frac{1}{\Delta x}\left(g_a.W_{xx} + g_t.\left(W^{t1}_{xx} + W^{t2}_{xx}\right)\right) \tag{77}$$

And so on for all equations (65-76).

If implementation is based on three eigenvectors (e.g, from DTI), then the orientation components of the two transverse vectors directly determine $W_{ei}^{t1}$ and $W_{ei}^{t2}$ for each direction through the same equations (47-63) as the primary orientation vector (**Fig. S3**). However, if only information on the primary eigenvector is defined, then we must define the transverse vectors from this. There are an infinite set of vectors orthogonal to the primary eigenvector, so the choice is somewhat arbitrary.

The most straightforward approach is to exploit the angle between the x-y plane and the z-axis (which we will call ϕ, as it is an elevation angle), which enables a π/2 rad rotation in the plane defined by the orientation vector and the z-axis to be trivially implemented (**Fig. S3**). The first set of transverse orientation components can therefore be described by:

$$\phi = \sin^{-1} Oz \tag{78}$$

$$Ox_{t1} = \left(Ox/\cos(\phi)\right).\cos(\phi + \tfrac{\pi}{2}) \tag{79}$$

$$Oy_{t1} = \left(Oy/\cos(\phi)\right).\cos(\phi + \tfrac{\pi}{2}) \tag{80}$$

$$Oz_{t1} = \sin(\phi + \tfrac{\pi}{2}) \tag{81}$$

Note the ϕ defined here is not the conventional, spherical polar coordinate ϕ, but, rather, represents an angle measured anti-clockwise from the x-y plane. We can then define a vector orthogonal to both the primary eigenvector and this defined transverse eigenvector by their cross product, giving the second transverse vector as (**Fig. S3**):

$$Ox_{t2} = Oy.Oz_{t1} - Oz.Oy_{t1} \tag{82}$$

$$Oy_{t2} = Oz.Ox_{t1} - Ox.Oz_{t1} \tag{83}$$

$$Oz_{t2} = Ox.Oy_{t1} - Oy.Ox_{t1} \tag{84}$$

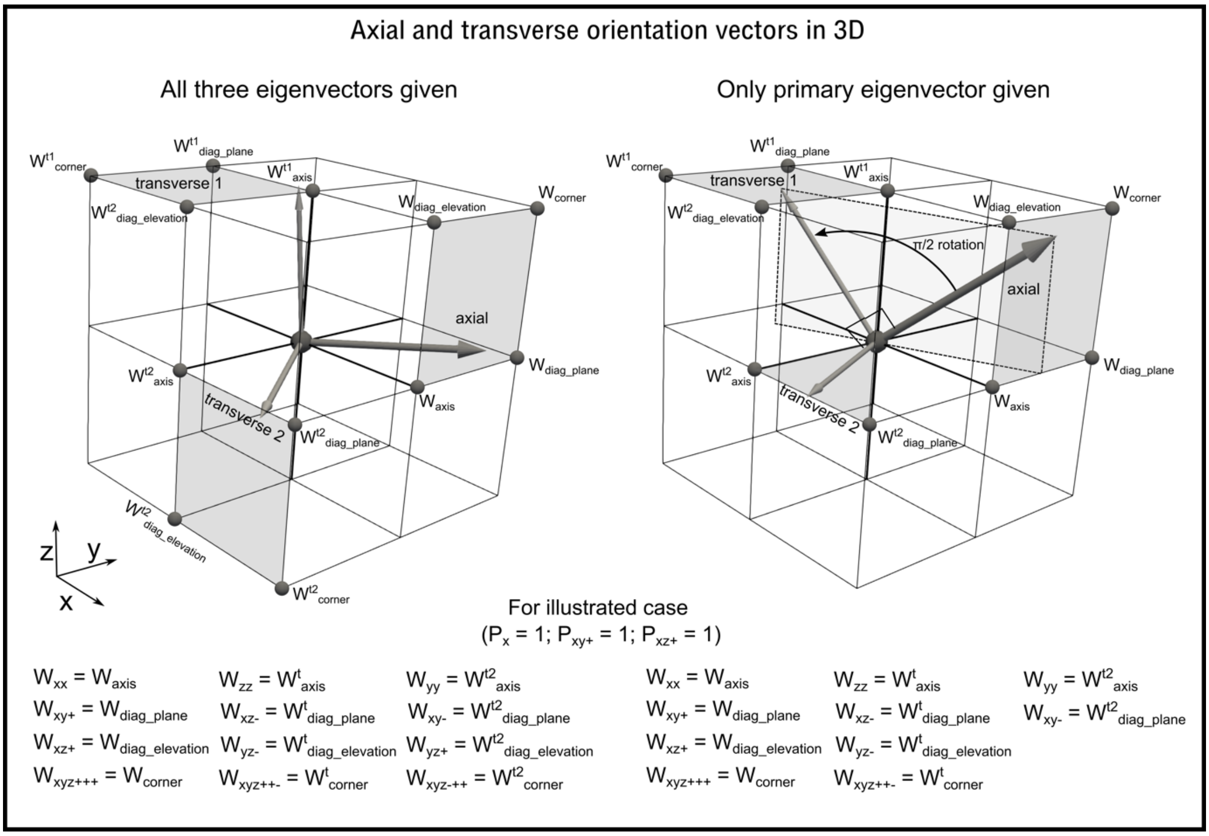


**Figure S3.** Illustration of the transverse orientation vectors and the directions to which they contribute in 3D. **Left** - illustration of the three orientation vectors as given by imaging data; each vector points in a quadrant, contributing to up to four directions. **Right** – illustration of the definition of the two transverse vectors if only the primary eigenvector is given. From the axial orientation, a π/2 rotation is applied towards the z-axis to define transverse vector 1. The cross-product of these two vectors then defines transverse vector 2, which will always point in the x-y plane. Note that only contributions in the positive direction are shown for clarity: all coupling is in both directions along each vector, as illustrated in **Fig. S2B.**

This has the implication that the vector corresponding to the first transverse direction will always be in the plane defined by the primary eigenvector and the z-axis, and the second transverse direction will always be in the x-y plane **Fig. S3**). These are arbitrary choices but provide a simple, convenient and consistent approach to defining these vectors when the structural data do not contain this information. Then, $W_{ei}^{t1}$ and $W_{ei}^{t2}$ for each direction is determined by passing the new vector components through the same equations as the primary vector.

This has a special case where z = 1 ($\phi = \pi/2$ rad and $\theta_{xy}$ is undefined), in which case it is trivial and consistent to apply the transverse directions to the x- and y-axis directions, by defining transverse vector 1 as (1,0,0) and transverse vector 2 as (0,1,0).

#### 1.4 Cellular dynamics and model parameters

Example simulations presented in this study were performed using cellular electrophysiology described by the hybrid minimal model presented in Colman (2019). This is a simple model parameterised for the magnitudes of ionic currents and voltage range observed in cardiac electrical excitation.

It is useful and desirable to be able to define the connection parameter conductances ($g_a$ and $g_t$) from the diffusion coefficient, $D$, to enable easy implementation to replace FDM, or similar, where $D$ has been defined to match conduction velocity. A conversion factor can be obtained through analysis of a simple case in 2D where myocyte orientation is in the x-direction, $D$ is homogeneous, and the spatial step is equal in each direction. The FDM discretisation in this case reduces to (Benson et al., 2021):

$$\nabla\left(\mathbf{D}\nabla V\right)\approx D1\frac{\Delta Vx}{\Delta x^2}+D2\frac{\Delta Vy}{\Delta x^2} \tag{85}$$

Which, implementing the central differences approach (Benson et al., 2021), becomes:

$$\nabla\left(\mathbf{D}\nabla V\right)\approx D1\frac{\left(V^{x+1,y}+V^{x-1,y}-2V^{x,y}\right)}{\Delta x^2}+D2\frac{\left(V^{x,y+1}+V^{x,y-1}-2V^{x,y}\right)}{\Delta x^2} \tag{86}$$

In this condition, the network model, where node $i = x,y$, reduces to:

$$\sum_n I_{gap}^{(i,n)} = \frac{g_a\left(V^{x+1,y}-V^{x,y}\right)}{\Delta x}-\frac{g_a\left(V^{x,y}-V^{x-1,y}\right)}{\Delta x}+\frac{g_t\left(V^{x,y+1}-V^{x,y}\right)}{\Delta x}-\frac{g_t\left(V^{x,y}-V^{x,y-1}\right)}{\Delta x} \tag{87}$$

Which simplifies to:

$$\sum_n I_{gap}^{(i,n)} = g_a\frac{\left(V^{x+1,y}+V^{x-1,y}-2V^{x,y}\right)}{\Delta x}+g_t\frac{\left(V^{x,y+1}+V^{x,y-1}-2V^{x,y}\right)}{\Delta x} \tag{88}$$

Comparing these equations yields the conversion factor of 1/Δx:

$$g_a = \frac{D1}{\Delta x} \tag{89}$$

$$g_t = \frac{D2}{\Delta x} \tag{90}$$

As will be shown in results, this conversion gives the same conduction velocities as the FDM method in these simple conditions. Note that this linear relationship means $g_t$ can be defined from the anisotropy ratio (AR) and $g_a$, in the same way as often for D1 and D2:

$$g_t = g_a / AR \tag{91}$$

In this setup, the units of $g_a$ and $g_t$ are the same as those for the maximal conductances of the ion currents (nS/pF), as the membrane capacitance ($C_m$) in equation (1) has been absorbed into these maximal conductance terms (equation 2). However, this model does not require this to be the case and enables coupling between cells of different $C_m$ while still conserving current; one could define $g_a$ as an absolute conductance (nS) and maintain inclusion of $C_m^{-1}$ when implementing equation (1).

##### 1.4.1 Computational efficiency

The simplest computational implementation of this approach would be to determine the gap junctional currents associated with each node (the sum in equations 1-2), expanding as illustrated in **Fig. S1B**. In FDM, the coupling between two nodes can contribute differently to each node (as each node's term depends on the local D and D gradients, and these may be different factors for two adjacent nodes); thus, this *requires* the coupling term to be calculated on a node-by-node basis. However, because this network approach implements a strategy that implicitly conserves current, wherein the exact same junctional current applies to both nodes comprising the junction (and the conductance of that junctional current is determined from *both* nodes), this simple implementation contains computational redundancies wherein the same gap junctional current is always calculated twice, once for each node associated with the junction. To improve computational efficiency, this method can be implemented in such a way that the currents for all individual and unique gap junctions are calculated within a single loop, with this junctional current contributing to the sum in equations (1-2) for each of the two nodes associated with that junction (identified by the junction maps, equations 11-12, and applied with the appropriate sign, equations 4-5). Practically, this can be more easily implemented by updating the voltage differential for the two nodes when each junctional current is calculated. This halves the number of calculations performed associated with the coupling term, compared to the simple approach or FDM (where each junction is indeed calculated twice), at improved computational efficiency.

### 2 Connection maps in 3D

Connection maps in each direction are shown for two 3D geometrical models: a human ventricular wedge (Benson et al., 2007) where only the myocyte orientation was given (and so the two transverse directions are calculated within the model; **Fig. S4**), and a full reconstruction of rat bi-ventricular geometry (Whittaker et al., 2019) wherein all three orientations describing cardiac tissue structure (myocyte, sheet and sheet normal, corresponding to the three DT-MRI eigenvectors) were given (**Fig. S5**).

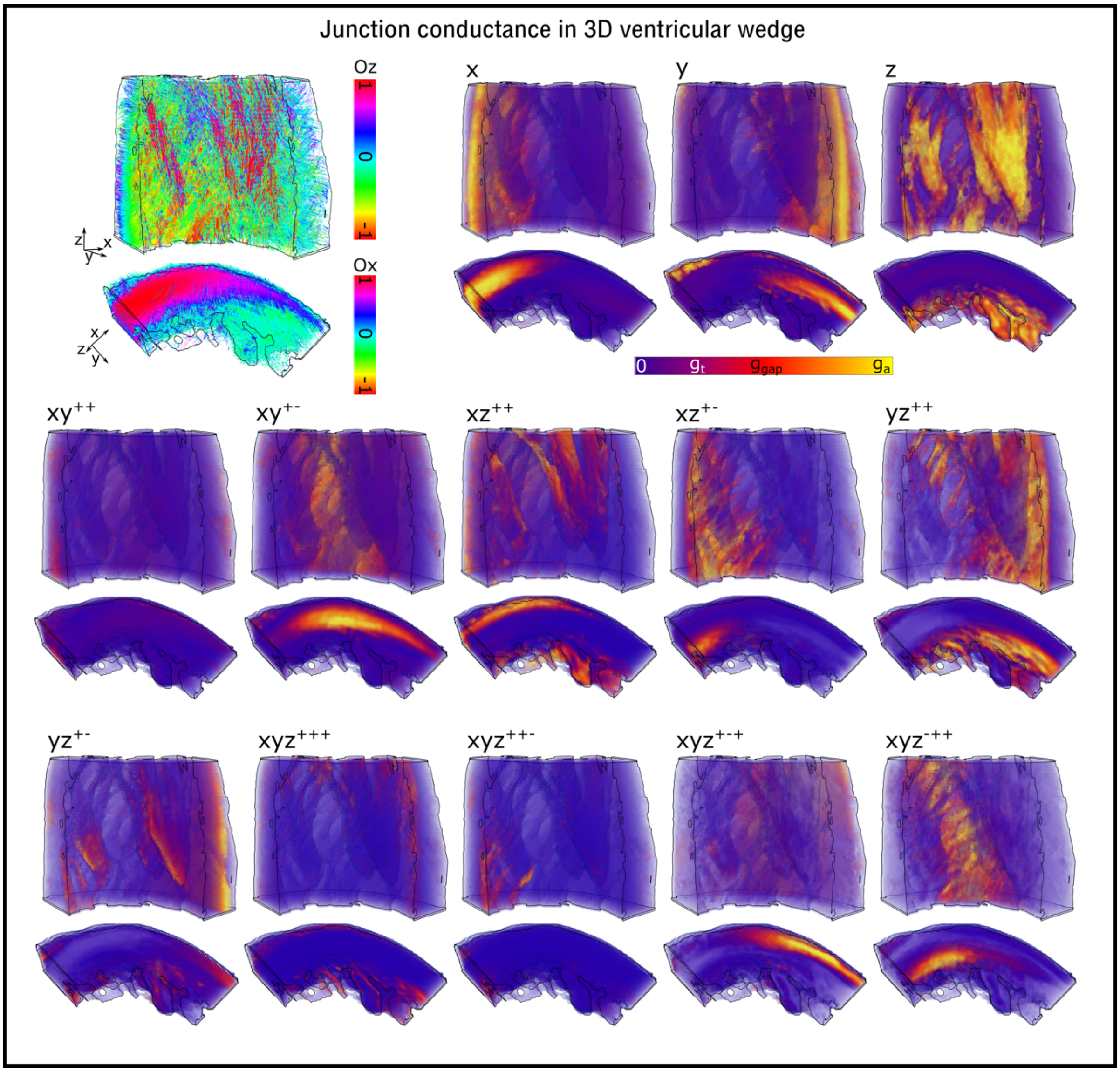


**Figure S4**. Illustration of directional condutances in a 3D human ventricular wedge model. Myocyte orientation streamlines are shown to provide context, with colour corresponding to either the z- or x-component (dependent on the view), along with the magnitude of the connection for each direction. The anisotropy ratio is 4:1 (i.e., $g_a = 4g_t$). As with **Fig S4**, for clarity of visualisation, the junctional conductances defined in equations (64-76) are not scaled by the $1/\Delta x$ etc factors, in order to normalise between axis and diagonal conductances.

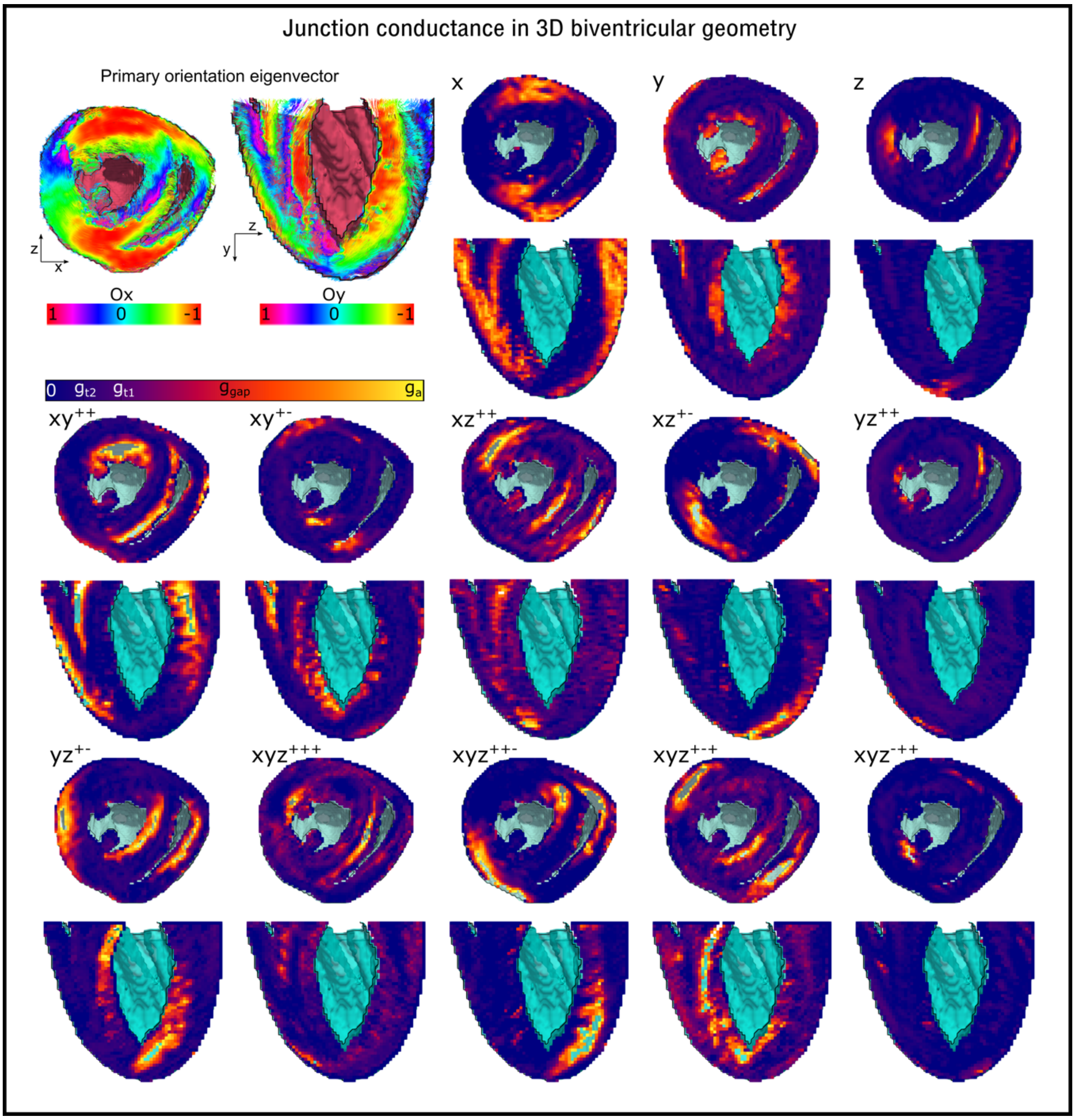


**Figure S5**. Illustration of directional conductances in a full bi-ventricular model, with all three eigenvectors given. Primary myocyte orientation streamlines are shown to provide context, with colour corresponding to either the x- or y-component (dependent on the view), along with the magnitude of the connection for each direction. Orientation and conductances are visualised on two slices, rather than throughout the volume, for clarity. The anisotropy ratio is 8:2:1 (i.e., $g_a = 4g_{t1} = 8g_{t2}$). As with **Fig. S4**, for clarity of visualisation, the junctional conductances defined in equations (64-76) are not scaled by the $1/\Delta x$ etc factors, in order to normalise between axis and diagonal conductances.

#### 2.1 Connection maps from distributions

Examples of connection maps where connection strength is sampled from defined distributions are show in **Fig. S6**.

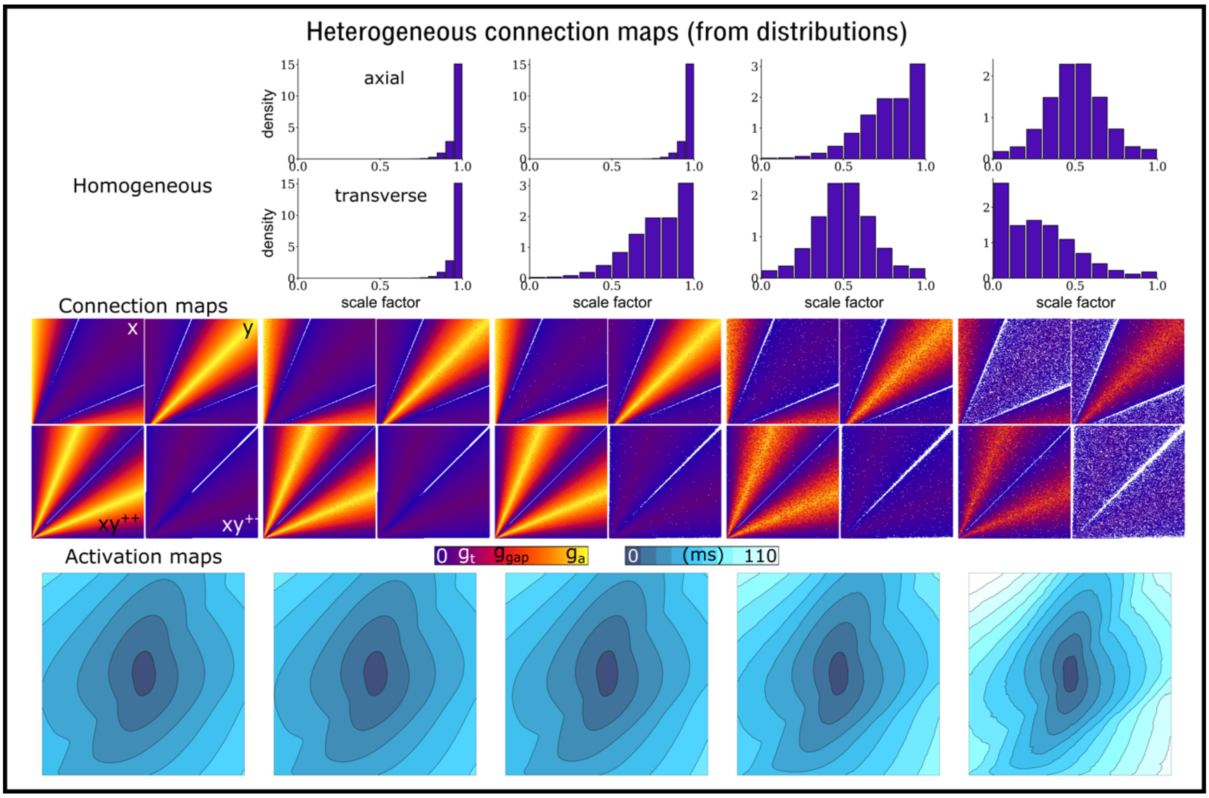


**Figure S6**. Connection maps and activation patterns associated with continuously distributed connection heterogeneity. Top panels show frequency density histograms of the scale factor used to modify network connections (axial - upper; transverse – lower), illustrating different distributions. Middle panels illustrate the connection maps associated with each set of distributions, and the bottom panel illustrates activation patterns resulting from these heterogeneous media.

### 3 Symmetry considerations

Both the FDM and network model do not give completely symmetric behaviour: when myocyte orientation is in the diagonal, the longitudinal and transverse conduction velocities are not exactly equal to that if the orientation points along an axes (**Fig. S7**). This is a consequence of the reaction term and role of the rapid, threshold-based $I_{Na}$ upstroke in determining conduction velocity – this timing is equal whether coupled cells are along an axes or diagonal, independent of the increased distance between nodes. Due to the direct formulation of the network model, it is possible to introduce correction factors in order to maintain this symmetry, if desired (the asymmetry is an acceptable approximation, so this is not an absolute necessity). By introducing empirically derived factors which multiply the axial and transverse connections in the diagonal and corner directions (but not the main axes directions) individually, symmetry may be restored (**Fig. S7**).

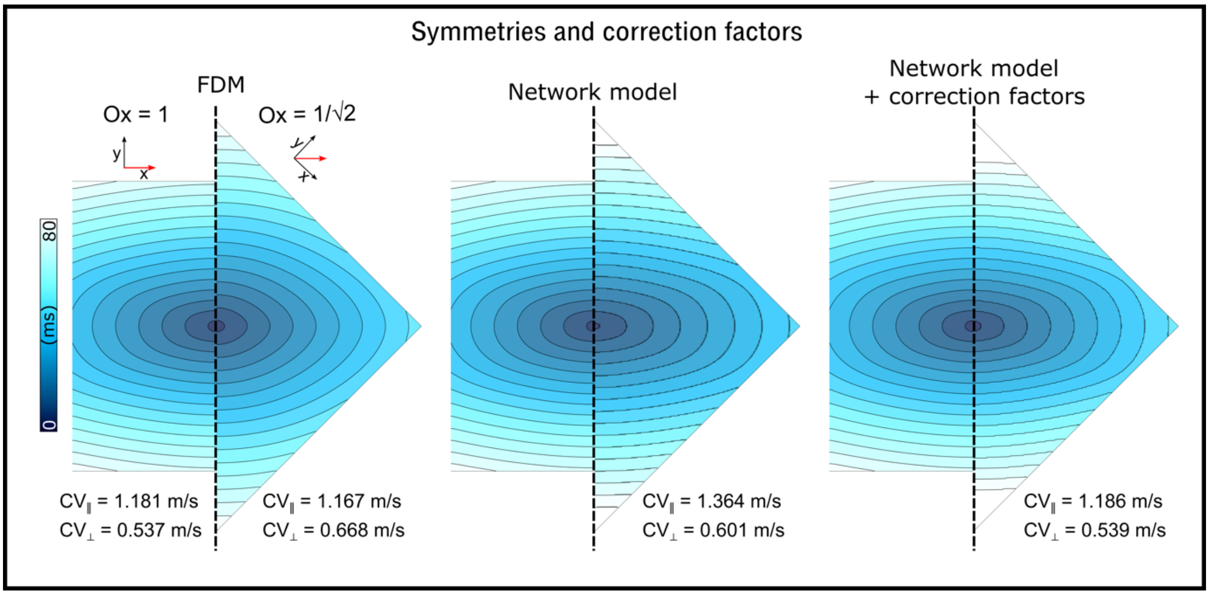


**Figure S7**. Symmetry considerations in FDM and the network models. Two simulations are compared for each model, with the orientation pointing in either the x or xy+ directions. Half of the tissue is shown for each condition, with the xy+ condition rotated by π/4 to enable direct comparison of the activation contours. The correction factors used to match the symmetry were 0.77 and 0.85 on the diagonals, for the axial and transverse terms, respectively.

*Note: this whole effect is emphasised by the minimal model, which has a large* $I_{Na}$ *and rapid upstroke; in other models tested, the effect is substantially smaller.*
